# Integrated metagenome-resolved profiling of the resistome, virulome, and mobilome in the gut microbiota of wild birds

**DOI:** 10.1101/2025.07.09.663844

**Authors:** Jin-Wen Su, Ying-Qian Gao, Hany M. Elsheikha, Cong-Cong Lei, Hong-Lin Xie, Fu-Long Nan, He Ma, Meng-Ting Yang, Hai-Tao Wang, Hong-Bo Ni, He-Ting Sun, Hong-Chao Sun, Xiao-Xuan Zhang

## Abstract

Wild birds, with their extensive geographic distributions and high mobility, are increasingly recognized as important players in the dissemination of antimicrobial resistance. Their gut microbiota, shaped by exposure to diverse environments, may act as both reservoirs and vectors of antibiotic resistance genes (ARGs), virulence factor genes (VFGs), and mobile genetic elements (MGEs). In this study, we reconstructed 2,516 high-quality metagenome-assembled genomes (MAGs) from 718 gut metagenomes of wild birds to comprehensively profile their resistome and virulome. We identified 5,596 ARG-encoding proteins across 389 distinct ARG types, with multidrug resistance emerging as the most dominant category. *Escherichia coli* was the principal carrier of ARGs, and genes conferring resistance to elfamycin antibiotics via target alteration were notably widespread—indicating persistent antibiotic selection pressures in avian habitats. Co-occurrence analyses revealed extensive genetic linkage between ARGs, VFGs, and MGEs. Critically, we detected 25 ARG–MGE co-localization events within 5-kilobase genomic regions, highlighting a strong potential for horizontal gene transfer and accelerated resistance dissemination within microbial communities. Of particular concern was the detection of the *tetX1* gene—conferring resistance to tigecycline, a last-resort antibiotic—in the gut microbiota of *Chroicocephalus ridibundus* and *Cygnus cygnus*. This finding strongly implicates anthropogenic pollution in the spread of clinically relevant ARGs into wildlife and emphasizes the risk of environmental transmission to other hosts, including humans. These results underscore the critical ecological role of wild birds in the global antimicrobial resistance network. As both reservoirs and potential vectors of ARGs, they represent a significant but under-monitored interface between environmental and clinical resistance pathways. Enhanced surveillance and mitigation strategies targeting wildlife are urgently needed to curb the environmental propagation of antimicrobial resistance.

## Introduction

Wild birds inhabit an array of ecosystems—from high-altitude lakes and coastal plains to marine environments—requiring exceptional ecological adaptability. This adaptability extends to their gut microbiota, which co-evolves with the host to form complex microbial communities suited to varied habitats. These microbiomes perform essential functions, including nutrient metabolism and immune modulation, while the host provides a stable niche and resources. The composition and function of the gut microbiota are strongly influenced by host species and their specific environmental contexts, shaping a dynamic equilibrium between symbiosis and pathogenic potential ^1–3^.

Due to their global distribution and migratory behavior, wild birds play a pivotal role in the long-range dissemination of infectious agents. Their ability to traverse vast distances creates a conduit for the intercontinental spread of pathogens, including antibiotic-resistant bacteria (ARB) ^4,5^. Studies have shown that wild birds residing near anthropogenic environments, such as urban or agricultural areas, tend to harbor bacterial strains with higher levels of antibiotic resistance compared to those from remote habitats ^6^. The first recorded instance of ARB in wild birds dates back to the 1970s, when pigeons were found to carry multidrug-resistant strains, including resistance to chloramphenicol ^7^. Since then, growing evidence has confirmed that migratory birds act as vectors for ARB across ecosystems ^8^. For instance, resistant *Escherichia coli* strains isolated from black-headed gulls in Sweden were genetically indistinguishable from those found in human clinical samples ^9^. This indicates a bidirectional transmission potential between wildlife and humans, particularly in areas with high environmental contamination ^10^.

Molecular typing methods, such as ERIC-PCR, have revealed closely related *E. coli* clones among humans, poultry, and wild birds, suggesting shared reservoirs and gene flow across species boundaries ^11^. The spread of ARB is driven by various factors, including unregulated antibiotic use, inadequate sanitation, and environmental exposure to human and animal waste. Consequently, wild birds are not only sentinels but also potential amplifiers of resistance dissemination.

The gut microbiota of wild birds may carry and transmit antibiotic resistance genes (ARGs), posing significant implications for public health ^12^. Understanding the structure and function of these microbial communities—and their associated ARGs, virulence factor genes (VFGs), and mobile genetic elements (MGEs)—is essential for mitigating the environmental spread of resistance. ARGs enable microorganisms to withstand antimicrobial agents, while VFGs enhance pathogenicity by promoting host invasion and immune evasion ^13^. MGEs such as plasmids, transposons, and bacteriophages facilitate the horizontal transfer of these genes, accelerating the evolution and dissemination of both resistance and virulence traits ^14,15^.

This study investigates the resistome and virulome of the wild bird gut microbiota using a large-scale metagenomic dataset. Specifically, we profile the diversity and distribution of ARGs, VFGs, and MGEs, and explore their co-occurrence and potential for horizontal gene transfer. Our findings provide novel insights into the ecological role of wild birds in antimicrobial resistance dissemination and highlight the importance of integrated surveillance strategies to safeguard public and environmental health.

## Result

### Comprehensive genomes of gut microbiota from wild birds

We analyzed 718 gut metagenomic samples collected from wild birds encompassing 16 orders, 29 families, 43 genera, and 91 species (Supplementary Table 1). Assembly and binning of the metagenomic data yielded a total of 10,455 metagenome-assembled genomes (MAGs). Following stringent quality filtering (completeness >50% and contamination <10%), 3,181 high-quality MAGs were retained. These genomes were subsequently clustered at 99% average nucleotide identity (ANI), resulting in 2,516 non-redundant, strain-level representative MAGs (Fig. 1).

**Fig. 1:**
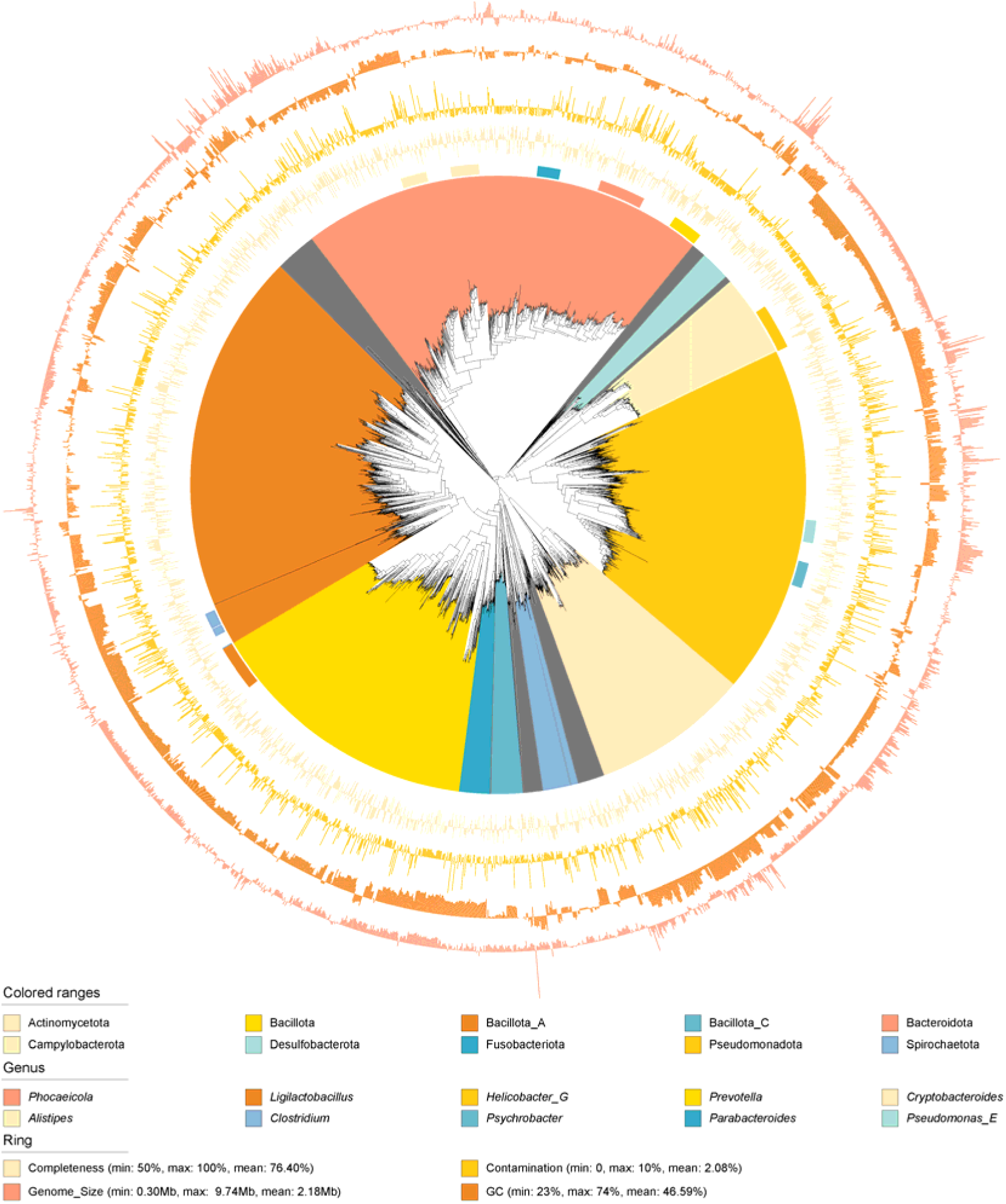
Phylogenetic Tree of 2,516 MAGs. The branches are color-coded according to phylum-level classification, with the first outer ring representing genus-level classification. The second outer ring shows a bar chart illustrating the completeness of each genome, while the third outer ring indicates the level of contamination. The fourth outer ring represents genome size, and the fifth outer ring depicts the GC content of the genomes.

The sizes of the representative MAGs ranged from 0.30 Mb to 9.74 Mb, with a mean size of 2.18 Mb. Genome completeness varied between 50% and 100% (mean: 76.40%), while contamination levels ranged from 0% to 10% (mean: 2.08%). The GC content of these genomes spanned 23% to 74%, with a mean of 46.59% (Fig. 1 and Supplementary Table 2).

Taxonomic annotation revealed that 22 MAGs belonged to the domain Archaea, while the remaining were classified as Bacteria. Overall, the dataset encompassed 34 phyla, 58 classes, 120 orders, 242 families, 658 genera, and 505 species. Notably, 1,720 MAGs (68.3%) could not be confidently assigned to any known species, indicating a high proportion of previously uncharacterized microbial diversity (Supplementary Table 2).

At the phylum level, the most abundant groups were *Bacteroidota* (21.22%), *Bacillota_A* (20.99%), *Pseudomonadota* (18.20%), and *Bacillota* (14.42%). At the genus level, the most prevalent taxa included *Phocaeicola* (2.35%), *Ligilactobacillus* (2.31%), *Helicobacter_G* (2.27%), and *Prevotella* (1.55%).

### The composition of ARGs in the Gut microbiota of wild birds

To characterize the antibiotic resistance gene (ARG) landscape in the gut microbiota of wild birds, we compared the 2,516 representative MAGs to the Comprehensive Antibiotic Resistance Database (CARD). This analysis identified 5,596 putative ARG-encoding proteins, representing 389 unique ARG types (Supplementary Table 3).

Among the identified ARGs, those conferring resistance to multiple antibiotic classes were the most prevalent, comprising 47.3% of the total, followed by ARGs associated with resistance to peptide antibiotics (7.2%), tetracyclines (5.66%), and aminoglycosides (5.4%) (Supplementary Fig. 1A). Mechanistically, the majority of ARGs functioned via four primary resistance mechanisms: antibiotic efflux (29.56%), antibiotic inactivation (27.25%), target alteration (26.48%), and multi-mechanism pathways (9%) (Supplementary Fig. 1B). At the abundance level, multi-drug resistance ARGs remained dominant (43.21%), with substantial contributions from ARGs conferring resistance to elfamycin (15.43%), aminoglycoside (7.15%), and fluoroquinolone (6.72%) antibiotics (Fig. 2A). The most common resistance mechanisms by abundance were target alteration (44.79%), efflux (38.55%), multi-mechanism (8.61%), and inactivation (3.77%) (Fig. 2B).

**Fig. 2:**
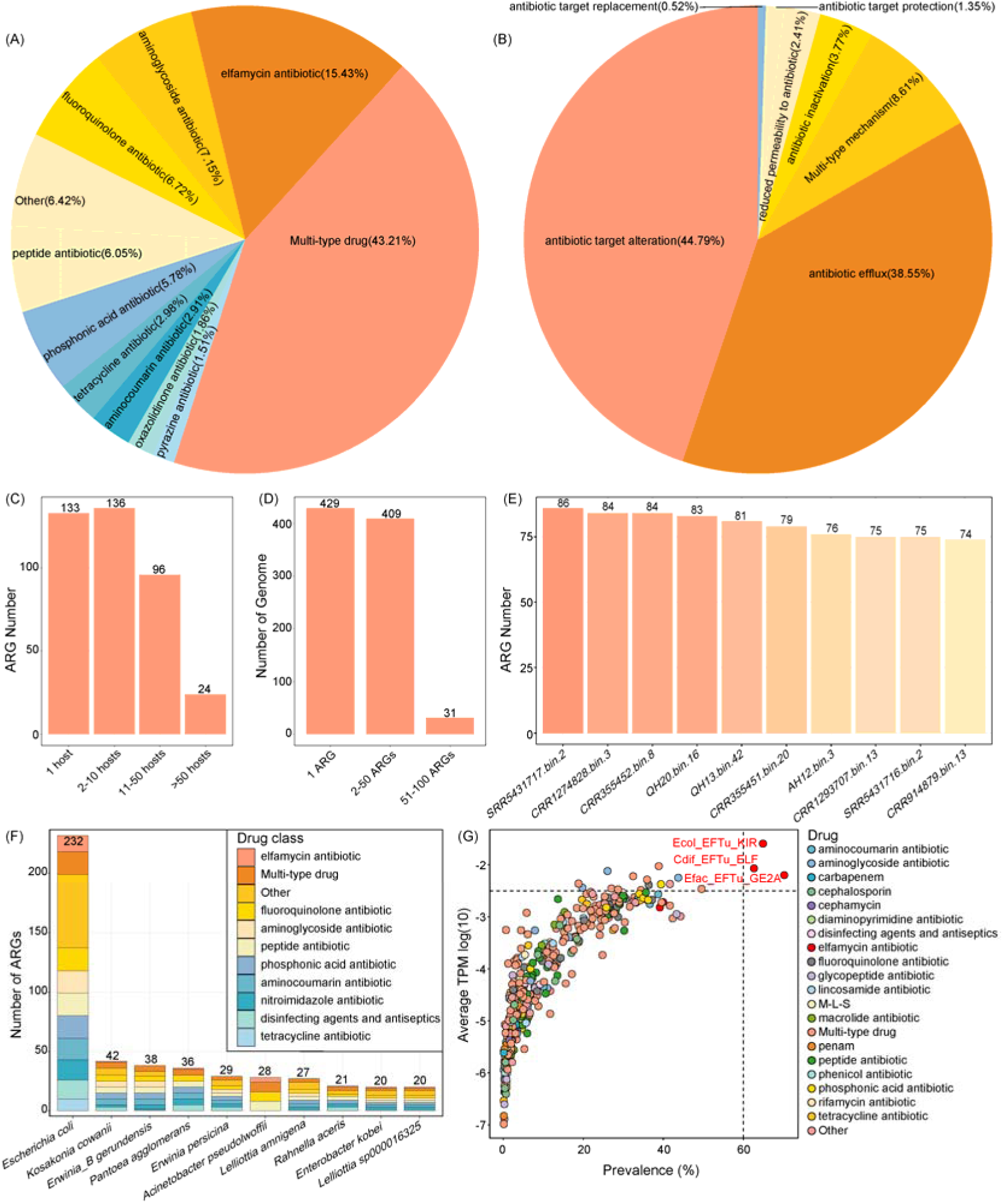
Distribution characteristics of ARGs in the gut microbiota of wild birds. (A) Distribution of the ten most abundant resistance phenotypes. (B) Distribution of resistance mechanism abundances. (C) Number of host species harboring each ARG. (D) Number of ARGs in each MAG. (E) MAGs with the top ten highest numbers of ARGs. (F) The ten bacterial species with the highest number of ARGs, with stacked colors representing corresponding resistance phenotypes. (G) Prevalence distribution of ARGs, with each point representing an ARG and its color indicating the resistance phenotype.

Distribution analysis of the 389 ARG types revealed that 133 (34.19%) were restricted to a single host species, 136 (34.96%) occurred in 2–10 hosts, 96 (24.68%) in 11–50 hosts, and 24 (6.17%) were widely distributed across more than 50 hosts (Fig. 2C). Of the 2,516 MAGs analyzed, 869 (34.54%) harbored at least one ARG. Specifically, 429 MAGs carried only one ARG, 409 MAGs carried between 2 and 50 ARGs, and 31 MAGs carried between 51 and 100 ARGs (Fig. 2D). The MAGs with the highest number of ARGs were *Escherichia coli* strains: SRR5431717.bin.2 (86 ARGs), CRR1274828.bin.3 (84), CRR355452.bin.8 (84), and QH20.bin.16 (83) (Fig. 2E). Overall, *E. coli* was the most frequent ARG carrier (232 MAGs), followed by *Kosakonia cowanii* (42), *Erwinia_B gerundensis* (38), and *Pantoea agglomerans* (36) (Fig. 2F).

The most prevalent ARG across the dataset was *Efac_EFTu_GE2A* (70.19%), followed by *Ecol_EFTu_KIR* (64.90%) and *Cdif_EFTu_ELF* (62.67%) (Fig. 2G). Notably, all three ARGs confer resistance to elfamycin antibiotics and operate via target alteration—a resistance mechanism indicative of antibiotic selection pressures in avian-associated environments (Supplementary Table 4).

Further correlation analysis within the top 20 most abundant bacterial families revealed that *Enterobacteriaceae*, *Mycobacteriaceae*, and *Xanthobacteraceae* were significantly associated with seven resistance phenotypes (R > 0.3, *p* < 0.001; Supplementary Fig. 2). These associations were primarily driven by key representative species, including *Escherichia coli* (linked to four resistance phenotypes), *Mycobacterium aubagnense* (two phenotypes), and *Pantoea deleyi* (two phenotypes) (R > 0.3, *p* < 0.01; Supplementary Fig. 3).

### The composition of VFGs in the gut microbiota of wild birds

To characterize the virulence potential of gut microbiota in wild birds, we analyzed 2,516 MAGs against the Virulence Factor Database (VFDB), identifying 8,090 open reading frames (ORFs) associated with virulence factor genes (VFGs). These ORFs were classified into 189 virulence factor (VF) types and grouped into 13 functional virulence factor categories (VFCs) (Supplementary Table 5).

The most abundant individual VFs by gene count were *Flagella* (28.11%), *Enterobactin* (4.85%), *Capsule* (4.82%), and *Polar flagella* (4.47%) (Supplementary Fig. 4A). At the functional category level, the dominant VFCs were associated with *Motility* (33.47%), *Adherence* (17.81%), *Effector delivery systems* (12.51%), and *Nutritional/Metabolic factors* (10.36%) (Supplementary Fig. 4B). When analyzed by abundance, *Flagella* remained the most prevalent VF (22.28%), followed by *Type VI secretion system (T6SS)* (8.18%), *Enterobactin* (7.91%), and *Elongation factor Tu (EF-Tu)* (6.33%) (Fig. 3A). Correspondingly, the top VFCs by abundance were *Motility* (23.18%), *Adherence* (19.44%), *Effector delivery systems* (18.94%), and *Nutritional/Metabolic factors* (14.46%) (Fig. 3B).

**Fig. 3:**
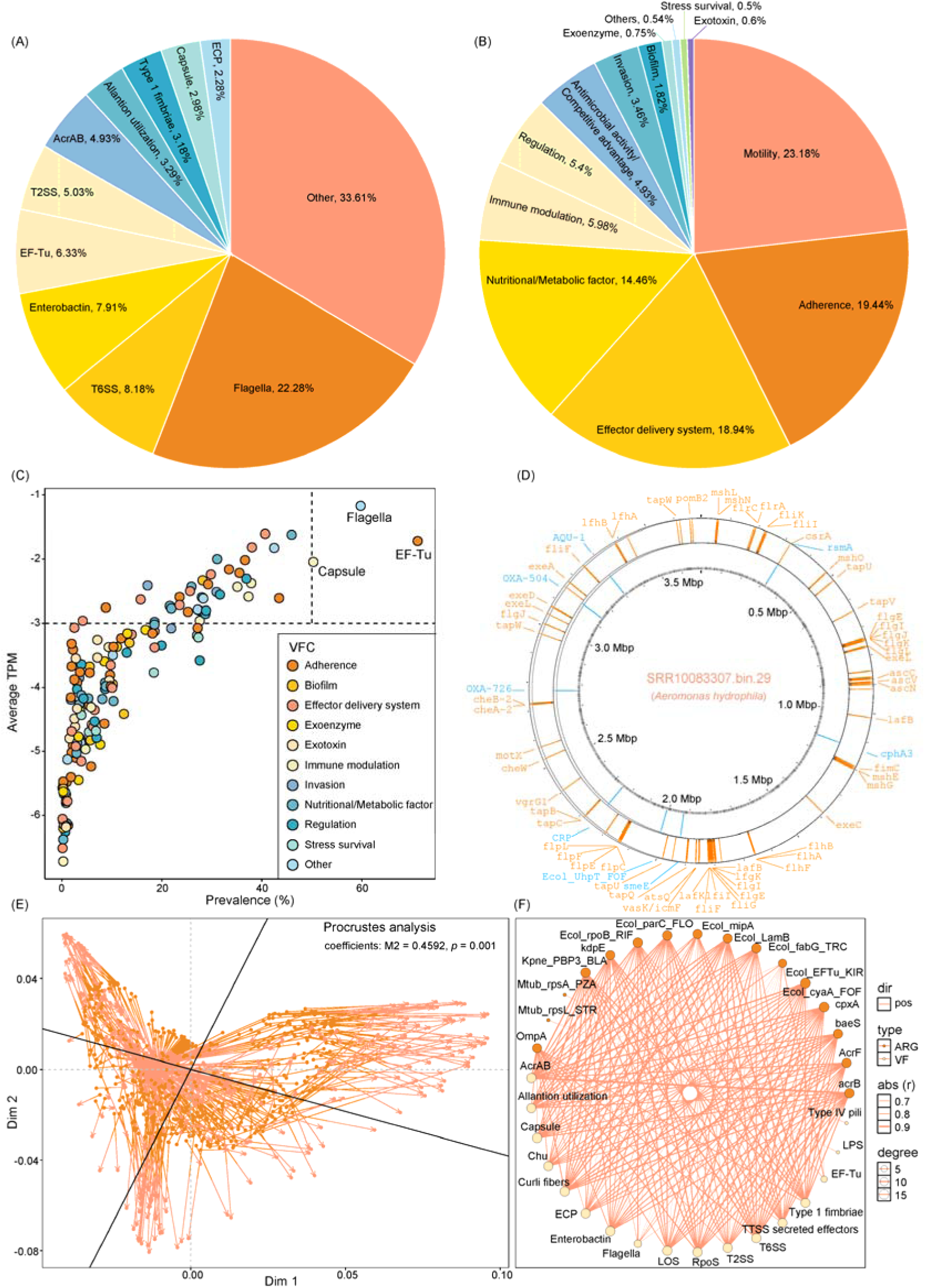
Distribution characteristics of VFGs in the gut microbiota of wild birds. (A) Distribution of the ten most abundant VFs. (B) Distribution of the ten most abundant VFCs. (C) Prevalence of VFs, with each dot representing a VF and the color indicating its corresponding VFC. (D) Genomic cycle map of SRR10083307.bin.29, with blue bars representing ARGs and yellow bars representing VFGs. (E) Procrustes analysis showing the correlation between the abundance profiles of ARGs and VFGs. (F) Correlation network between the top 20 most prevalent VFs and ARGs, based on Spearman’s rank correlation. Only significant associations with r > 0.6 and *P* < 0.05 are shown.

Of the 2,516 MAGs, 761 (30.25%) harbored at least one VFG (Supplementary Table 5). Notably, the MAG SRR10083307.bin.29, taxonomically assigned to *Aeromonas hydrophila*, carried the highest number of VFGs (*n* = 185) (Supplementary Fig. 4C), and also encoded eight ARGs (Fig. 3D). Prevalence analysis showed that *EF-Tu* was the most widely distributed VF (71.17%), followed by *Flagella* (59.75%) and *Capsule* (50.28%) (Supplementary Table 6).

To explore potential functional linkages, we examined the relationship between ARGs and VFGs. The richness of VFGs was strongly and positively correlated with ARG richness across MAGs (R = 0.96, *p* < 0.001; Supplementary Fig. 4D). Procrustes analysis confirmed a significant congruence between the compositional patterns of VFGs and ARGs (M² = 0.4592, *p* = 0.001; Fig. 3E). Further, Spearman correlation analysis of the top 20 most prevalent VFs and ARGs identified 194 significantly associated VF–ARG pairs (48.5%; *p* < 0.05), revealing intricate co-occurrence networks (Fig. 3F). The most pronounced association was observed between *acrB* and *AcrAB* (r = 0.94, *p* < 0.001), underscoring potential functional synergy between virulence and resistance determinants (Supplementary Table 7).

### The composition of MGEs in the gut microbiota of wild birds

MGEs are key drivers of horizontal gene transfer (HGT), facilitating the dissemination of ARGs among microbial populations. To assess the diversity and abundance of MGEs within the gut microbiota of wild birds, we analyzed 2,516 MAGs using a curated MGE database. In total, we identified 606 MGE-associated genes, corresponding to 294 distinct MGE types (Supplementary Table 8).

The most frequently detected MGEs by occurrence were *755_IS26_KT334335.1* (2.81%), *1233_tnpA_JN208880.1* (2.48%), *712_tnpA_EU850412.1* (1.98%), and *1004_tnpA_KT779035.1* (1.82%) (Supplementary Fig. 5A). These MGEs were classified into 18 categories, with *transposases* representing the majority (64.03%), followed by insertion elements *IS91* (15.02%), *Tn916* elements (6.27%), and *integrases* (4.29%) (Supplementary Fig. 5B).

When analyzed by abundance, *299_tnpA_AH010657.3* was the most prevalent, accounting for 23.83% of total MGE abundance, followed by *829_IS91_MOFD01000166.1* (4.68%), *1874_tnpA_JX843238.1* (4.37%), and *2364_tnpA_CP000864.1* (4.36%) (Fig. 4A). At the MGE type level, *transposases* dominated (75.62%), with lesser contributions from *IS91* elements (10.27%), *tniA* (2.83%), and *Tn916* (2.32%) (Fig. 4B).

**Fig. 4:**
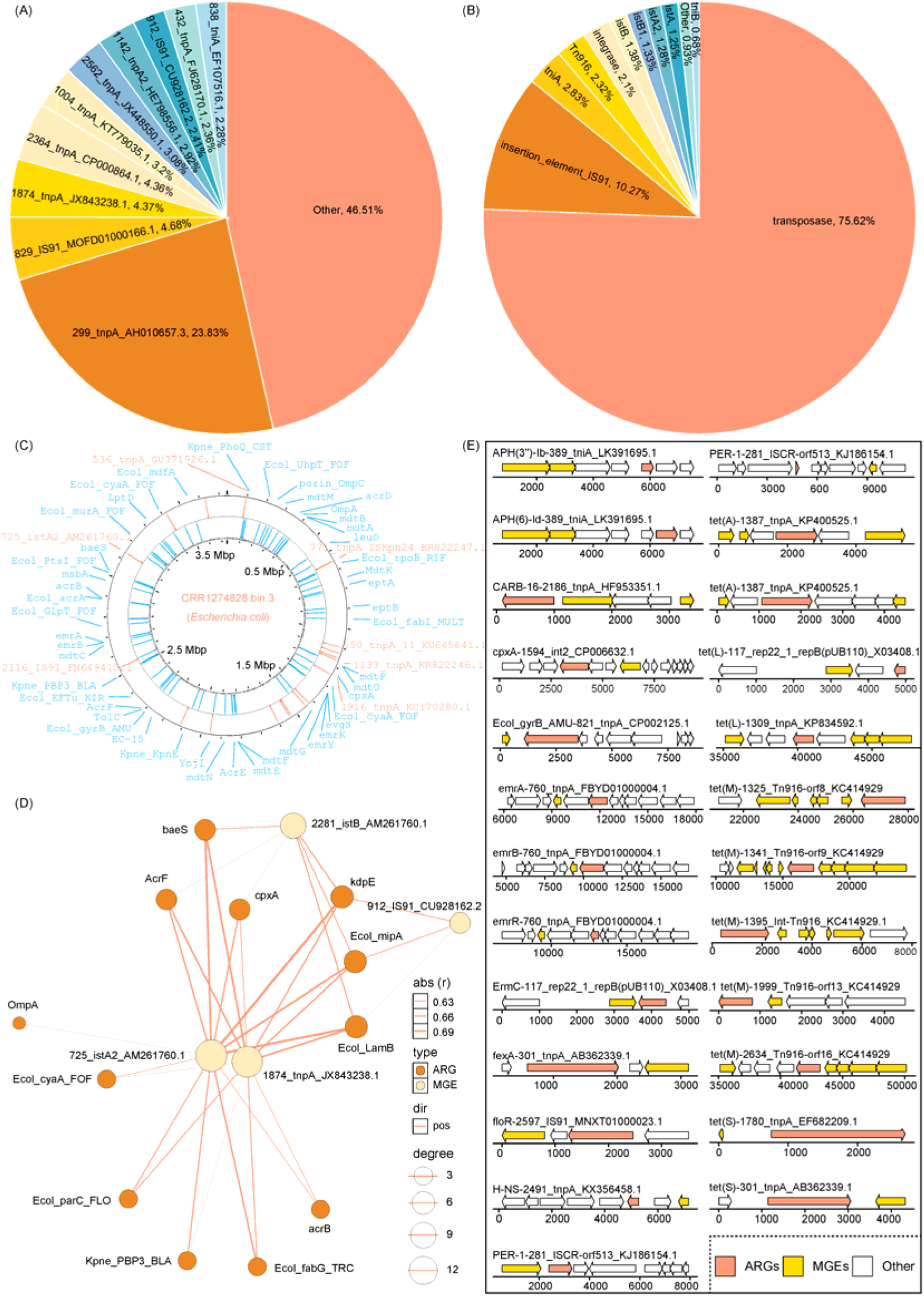
Distribution characteristics of MGEs in the gut microbiota of wild birds. (A) Distribution of the ten most abundant MGEs. (B) Distribution of the ten most abundant MGE types. (C) Genomic cycle map of CRR1274828.bin.3, with blue bars representing ARGs and yellow bars representing MGEs. (D) Correlation network between the top 20 most prevalent MGEs and ARGs, based on Spearman’s rank correlation. Only significant associations with r > 0.6 and *p* < 0.05 are shown. (E) Arrow diagram illustrating combinations of ARGs and MGEs within contigs. Right-pointing arrows represent genes on the forward DNA strand, while left-pointing arrows represent genes on the reverse strand.

Prevalence analysis revealed that *299_tnpA_AH010657.3* was the most widespread MGE (present in 45.13% of MAGs), followed by *287_tnpA_KT922275.1* (25.77%), *1004_tnpA_KT779035.1* (25.35%), and *1874_tnpA_JX843238.1* (24.37%) (Supplementary Fig. 5C; Supplementary Table 9). The MAG CRR1274828.bin.3, assigned to *Escherichia coli*, harbored the greatest number of MGEs (n = 23), and also encoded 84 ARGs, underscoring its high potential as a resistance gene reservoir (Supplementary Fig. 5D; Fig. 4C).

To examine the relationship between MGEs and ARGs, we perfo rmed correlation analyses. The richness indices of MGEs and ARGs were strongly and positively correlated (R = 0.87, *p* < 0.001; Supplementary Fig. 5E). Procrustes analysis confirmed a significant concordance between their abundance distributions across samples (M² = 0.7112, *p* = 0.001; Supplementary Fig. 5F). A correlation network constructed from the 20 most prevalent MGEs and ARGs revealed that *1874_tnpA_JX843238.1* and *725_istA2_AM261760.1* exhibited high centrality, each significantly associated with 11 and 12 ARGs, respectively (Fig. 4D; Supplementary Table 10).

To further elucidate potential HGT events, we identified ARGs located within ±5 kilobases (kb) of MGE sequences, suggesting physical linkage and potential mobility. This analysis revealed 25 unique MGE–ARG combinations, involving 18 distinct ARGs (Fig. 4E), including *tet(M)*, *tet(L)*, and *Ecol_gyrB_AMU*, among others. These proximally co-localized MGE–ARG pairs provide compelling evidence for the role of MGEs in mobilizing resistance determinants within wild bird-associated microbiota.

### Factors influencing the composition of ARGs in the gut microbiota of wild birds

To identify the factors shaping ARG composition in the gut microbiota of wild birds, we employed PERMANOVA based on the Bray-Curtis distance matrix. Marginal effect analysis revealed that bird species, sampling area, sampling time, and migratory status all significantly influenced ARG composition (*p* < 0.01; Fig. 5A). Among these, host species was the most influential factor, accounting for 30.22% of the variation. Additionally, multifactor analysis indicated a significant interaction between sampling area and host species in shaping ARG composition (Supplementary Fig. 6A).

**Fig. 5:**
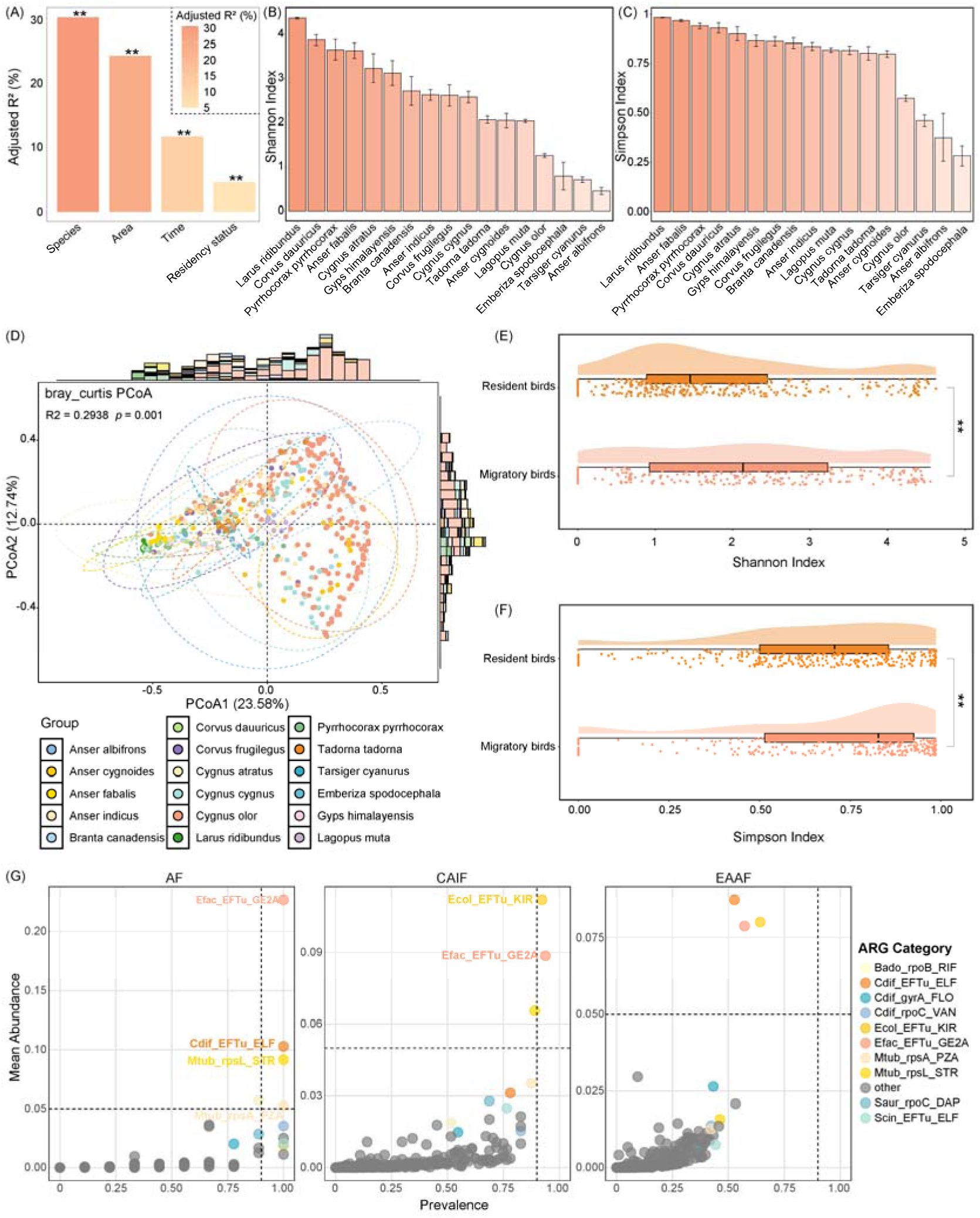
Analysis of factors influencing ARG composition. (A) PERMANOVA analysis based on Bray-Curtis distance matrix identifying key factors affecting ARG composition. The y-axis represents the adjusted R² values; ** denotes *P* < 0.01. (B-C) Shannon and Simpson indices of gut microbial ARGs across different bird species. Bars represent mean values, and error bars denote standard errors. (D) PCoA based on Bray-Curtis distances showing beta diversity differences in ARG composition among wild birds. Each point is plotted according to the first and second principal coordinates (PCoA1 and PCoA2). Ellipses represent the 95% confidence interval for each group, while stacked histograms on the top and right display sample density distributions. (E-F) Boxplots showing Shannon and Simpson indices of gut ARGs between resident and migratory birds. Statistical significance was assessed using the Wilcoxon rank-sum test: * *p* < 0.05; ** *p* < 0.01; *** *p* < 0.001; **** *p* < 0.0001. (G) Prevalence of gut microbial ARGs in wild birds across three major migratory flyways, with each dot representing an individual ARG and color indicating the ARG category.

Significant differences in ARG diversity were observed among bird species (Fig. 5B, 5C). *Larus ridibundus* exhibited the highest Shannon and Simpson diversity indices for ARGs in its gut microbiota. Pairwise comparisons revealed significant differences in the Shannon index for 75% of species pairs and in the Simpson index for 71.32% of pairs (Supplementary Table 11). Principal Coordinates Analysis (PCoA) further confirmed significant variation in ARG composition across species (R² = 0.2938, *p* = 0.001).

ARG diversity also exhibited significant variation across sampling areas (*p* < 0.05; Supplementary Fig. 6B, 6C, 6D). Samples from New Zealand and Japan showed notably higher ARG diversity compared to those from other regions (*p* < 0.05). Similarly, significant differences in ARG diversity were observed between sampling years (*p* < 0.05; Supplementary Fig. 7A, 7B, 7C), with samples from 2020–2023 displaying higher diversity than those from 2016–2019. Residency status (migratory vs. resident) also had a significant impact on ARG diversity, with migratory birds exhibiting higher diversity than resident birds (*p* < 0.05; Fig. 5E, 5F; Supplementary Fig. 7D).

Further analysis of migratory birds grouped by flyway route revealed significant differences in ARG diversity among the groups (Supplementary Fig. 7E, 7F, 7G). The Central Asian–Indian Flyway (CAIF) had the highest ARG diversity, significantly surpassing the East Asian–Australasian Flyway (EAAF) (*p* < 0.0001).

We also examined the prevalence of specific ARGs in the gut microbiota of wild birds across three flyways. In the Atlantic Flyway (AF), the ARGs *Efac_EFTu_GE2A*, *Cdif_EFTu_ELF*, *Mtub_rpsL_STR*, and *Mtub_rpsA_PZA* were present at 100% prevalence (Fig. 5G; Supplementary Table 12). In the CAIF, *Ecol_EFTu_KIR* (92.19%) and *Efac_EFTu_GE2A* (93.75%) were prevalent in over 90% of samples. In the East Asian–Australasian Flyway (EAAF), the most prevalent ARGs were *Ecol_EFTu_KIR* (64.18%), *Efac_EFTu_GE2A* (57.21%), and *Ecol_rpoB_RIF* (53.23%).

### Wild bird gut microbiota-specific ARGs

To assess the specificity of ARGs in the gut microbiota of wild birds, we compared 2,516 MAGs obtained from wild bird gut microbiota with previously reported gut genomes from humans (36,467 MAGs) and chickens (17,013 MAGs). Functional annotation revealed that *Ecol_EFTu_KIR* was the most prevalent ARG across all three groups (Fig. 6A–C).

**Fig. 6:**
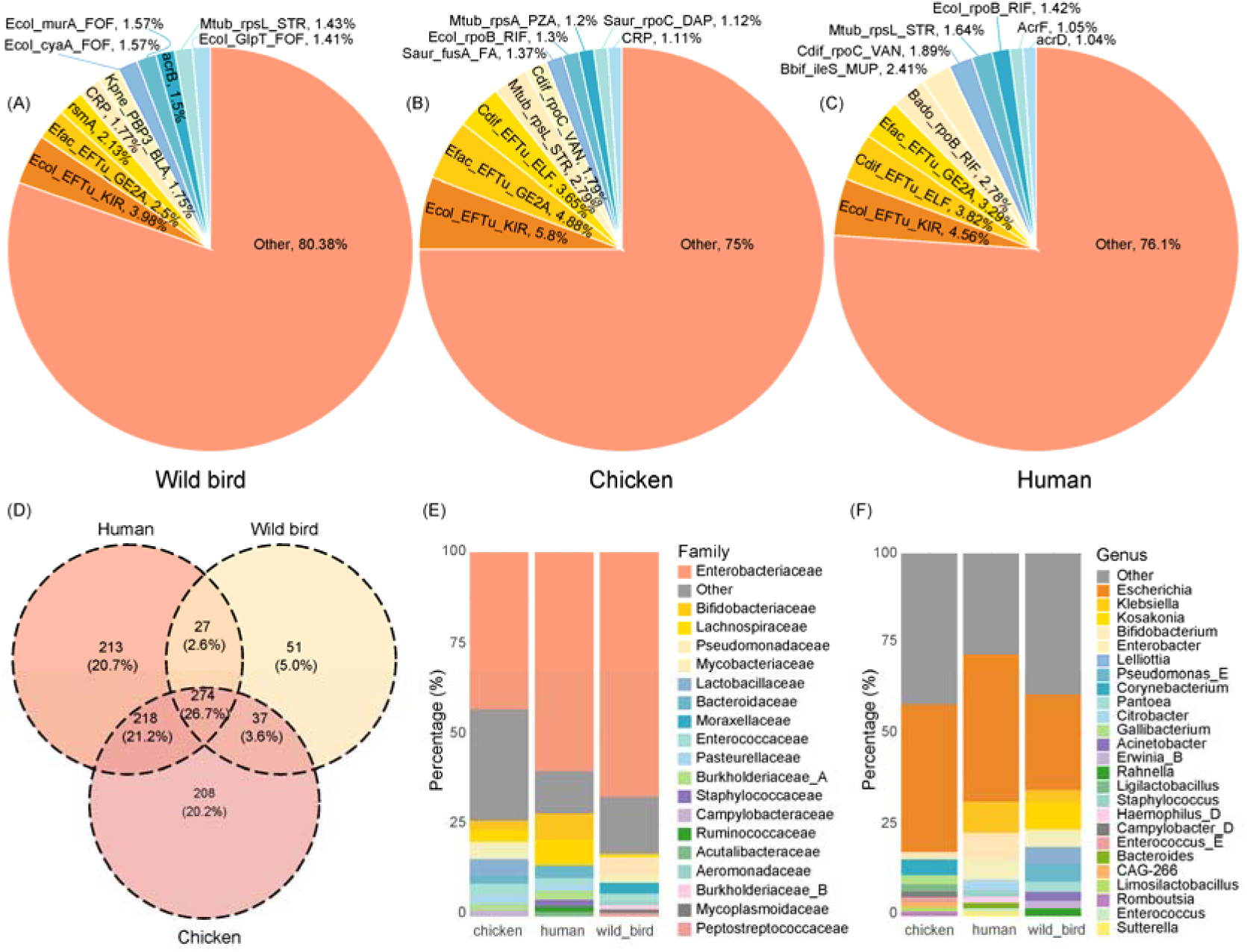
Comparison of ARG composition in the gut microbiota of wild birds, chickens, and humans. (A) Distribution of the ten most abundant ARGs in wild bird gut microbiota. (B) Distribution of the ten most abundant ARGs in chicken gut microbiota. (C) Distribution of the ten most abundant ARGs in human gut microbiota. (D) Venn diagram showing unique and shared ARGs among wild birds, chickens, and humans. (E) Taxonomic composition (family level) of microbial genomes carrying ARGs. (F) Taxonomic composition (genus level) of microbial genomes carrying ARGs.

In terms of resistance types, the dominant ARG classes in the gut microbiota of chickens and humans were multi-drug resistance, aminoglycoside antibiotic resistance, and peptide antibiotic resistance (Supplementary Fig. 8A, 8B). In contrast, the primary ARG types in the wild bird gut microbiota were multi-drug resistance, elfamycin antibiotic resistance, and aminoglycoside antibiotic resistance (Fig. 2A).

Regarding resistance mechanisms, the gut microbiota of chickens and humans primarily harbored ARGs associated with antibiotic inactivation, antibiotic efflux, and antibiotic target alteration, which collectively accounted for over 80% of ARGs (Supplementary Fig. 8C, 8D). In wild birds, antibiotic target alteration (44.79%) and antibiotic efflux (38.55%) were the most prevalent mechanisms (Fig. 2B).

Comparative analysis of ARG composition among the three host groups revealed 51 ARGs (5.0%) uniquely present in the wild bird gut microbiota (Fig. 6D). Additionally, 301 ARGs (29.3%) were shared with the human gut microbiota, and 311 ARGs (30.3%) were shared with the chicken gut microbiota. Among the ARGs shared with humans, 15 were identified as potentially mobile, including *tet(M)*, *rsmA*, *tet(L)*, *tet(S)*, *Ecol_gyrB_AMU*, *cpxA*, *emrB*, *emrA*, *emrR*, *H-NS*, *ErmC*, *APH(3’’)-Ib*, *APH(6)-Id*, *tet(A)*, and *floR* (Supplementary Table 13, Fig. 4E).

Additionally, we collected ARG sequences for four classes of drugs commonly used against Gram-negative bacteria: tigecycline, vancomycin, polymyxins, and β-lactams (Supplementary Table 14). Using stringent criteria (identity ≥90%, coverage ≥70%, E-value ≤1e-10), we conducted a BLASTn analysis to compare ARGs shared between wild bird and human gut microbiomes. Two genes met the criteria for tigecycline resistance (*tetX1*). These genes were identified in the gut microbiota of *Chroicocephalus ridibundus* and *Cygnus cygnus* and were taxonomically assigned to an unclassified species within the genus *Catellicoccus* and *Riemerella*, respectively.

## Discussion

Antibiotic resistance represents one of the most pressing global public health threats of the 21st century^16^. Wild birds, due to their extensive migratory patterns and frequent interactions with human-influenced environments, are emerging as critical vectors for the spread of ARGs ^17^. This study provides a comprehensive analysis of 718 gut microbial metagenomic samples from wild birds, representing 91 species across 29 families and 16 orders. By establishing a genomic resource, we reveal the composition of ARGs, VFs, and MGEs in wild bird gut microbiota, highlighting their co-occurrence patterns and shared ARG profiles with human and chicken microbiomes.

Our analysis identified 5,596 ARG-encoding proteins across 389 distinct ARG types, with multi-drug resistance genes comprising the majority (43.21%) of the ARG pool. This indicates that wild birds’ gut microbiota serves as a significant reservoir of ARGs. Notably, *Escherichia coli* was found to harbor the highest number of ARGs (232), consistent with previous studies that underscore its role as a major carrier of resistance genes ^18–20^. This finding underscores the importance of *E. coli* as a key player in the dissemination of antimicrobial resistance. Among the most prevalent ARGs in wild bird gut microbiota, several genes—such as *Efac_EFTu_GE2A*, *Ecol_EFTu_KIR*, and *Cdif_EFTu_ELF*—confer resistance to elfamycin-class antibiotics via a target alteration mechanism. These genes encode EF-Tu-related proteins, which serve as critical targets for protein synthesis and the primary binding site of elfamycin antibiotics ^21^. Structural modifications in EF-Tu reduce the binding affinity of these antibiotics, leading to resistance. The widespread occurrence of these target-altering ARGs suggests that specific selective pressures are shaping the microbial communities of wild birds’ environments ^22^.

VFG analysis revealed 189 types of VFs, with approximately 30.25% of MAGscontaining at least one VFG. The predominant VF types—flagella, EF-Tu, and T6SS—are closely associated with bacterial motility, host adhesion, and immune evasion ^23–25^. Notably, one *Aeromonas hydrophila* genome was found to harbor 185 VFGs and 8 ARGs simultaneously, indicating the presence of bacterial populations with both opportunistic pathogenicity and multi-drug resistance within wild bird guts. Further, significant positive correlations between VFGs and ARGs suggest that these factors may be co-selected under shared environmental pressures. For instance, the AcrAB-TolC efflux pump system has been shown to contribute to both antibiotic resistance and virulence regulation ^26^.

Our findings also highlight the co-localization of 25 ARG-MGE combinations within the wild bird gut microbiota, with ARGs located within a 5 kb region flanking MGEs. This spatial proximity supports the hypothesis that these resistance genes can spread across different microbial taxa through MGE-mediated horizontal gene transfer ^27^. For example, the co-localization of tetracycline resistance genes *tet(M)* and *tet(L)*—commonly associated with conjugative transposons like Tn916 and plasmids, respectively—demonstrates their potential for wide dissemination across bacterial species ^28–31^.

The composition of ARGs was significantly influenced by bird species, sampling region, sampling time, and residency status. Among these factors, host species emerged as the most influential, suggesting that species-specific physiological traits and ecological behaviors play a central role in ARG carriage ^32^. *Larus ridibundus* exhibited the highest ARG diversity, potentially reflecting a broader range of resistance genes in its gut microbiota ^33^. Analysis also revealed significant differences in ARG diversity among bird populations following different migratory flyways. These differences suggest that the environments traversed and the intensity of human activity along these routes shape the spatial distribution of ARGs ^32^. For example, migratory routes through regions with high human activity, such as agricultural discharges, may serve as hotspots for ARG dissemination. Therefore, targeted surveillance and intervention efforts along these flyways are urgently needed to mitigate the spread of resistance genes ^34–37^.

The study also identified 301 ARGs shared between wild birds and humans, and 311 shared with chickens, with 15 of these being potentially mobile ARGs. This highlights the risk of cross-species transmission ^38,39^. Of particular concern is the detection of the tigecycline resistance gene *tetX1* in the gut microbiomes of *Chroicocephalus ridibundus* and *Cygnus cygnus*, raising significant public health concerns. Tigecycline is considered a last-resort antibiotic for multidrug-resistant Gram-negative infections ^40^. The presence of its resistance gene in wild birds suggests that anthropogenic ARGs have infiltrated natural ecosystems via environmental pathways, amplifying the potential for long-distance transmission ^34,41^. Moreover, the detection of *tetX1* in an unclassified *Catellicoccus* species and *Riemerella anatipestifer* indicates potential intergeneric transmission ^42,43^, warranting further investigation into the mechanisms underlying its spread.

While this study provides valuable insights, it is not without limitations. The samples were unevenly distributed in terms of geographic coverage and species composition, and environmental variables such as habitat types and human activity intensity were not fully controlled. Future research, including long-term monitoring and experimental validation, will be essential to further unravel the dynamic evolution of ARGs and the factors influencing their spread in wild bird populations.

## Conclusion

This study analyzed 718 gut metagenomic samples from wild birds, reconstructing 2,516 high-quality MAGs to provide a comprehensive view of ARGs, VFGs, and mobile genetic elements (MGEs) within the gut microbiota of wild avian hosts. We identified 5,596 ARG-encoding proteins, representing 389 distinct ARG types, with multi-drug resistance being the most prevalent. *Escherichia coli* emerged as the primary carrier of ARGs, with several target modification-related genes conferring resistance to elfamycin antibiotics. Extensive co-occurrence among ARGs, VFGs, and MGEs was observed, with 25 ARG-MGE co-localization events identified, suggesting the potential for horizontal gene transfer and the synergistic evolution of resistance and virulence. Our findings also highlight that host species, sampling region, sampling time, and residency status all significantly influenced ARG diversity. Notably, birds migrating along the CAIF and AF flyways exhibited higher ARG burdens, pointing to the influence of environmental factors along migratory routes. Comparative analyses with human and chicken gut microbiomes revealed that wild birds not only harbor unique ARGs but also share potentially mobile resistance genes with other host groups. Of particular concern was the detection of the tigecycline resistance gene *tetX1* in *Larus ridibundus* and *Cygnus olor*, suggesting that anthropogenic contamination has infiltrated natural ecosystems, facilitating cross-boundary transmission of resistance genes. Our study underscores the critical role of wild birds as reservoirs and vectors of ARGs. These findings highlight the urgent need for cross-sectoral surveillance and further mechanistic research to mitigate the ecological and public health risks associated with antimicrobial resistance.

## Materials and methods

### Sample collection

A total of 50 cloacal fecal samples were collected from four wild bird species across four provinces in China (Supplementary Table 1). Cloacal fecal samples were obtained using sterile cotton swabs during fieldwork. Immediately after collection, the samples were stored at −20°C in portable freezers, transported to the laboratory, and then kept at −80°C until further processing. A subset of these samples was selected for metagenomic sequencing.

### Large-scale collection of wild bird gut microbiome metagenomic data

In addition to the samples collected in this study, 668 publicly available wild bird gut metagenomic samples were retrieved from several databases, including NCBI (https://www.ncbi.nlm.nih.gov/), ENA (https://www.ebi.ac.uk/ena/), CNCB (https://www.cncb.ac.cn/), and DDBJ (https://www.ddbj.nig.ac.jp/). Details of these samples can be found in Supplementary Table 1.

### Preprocessing of raw data

Raw reads were filtered for high quality using Fastp (v0.23.0) ^44^, followed by the removal of host genomic DNA using Bowtie2 (v2.5.0) ^45^. Contig assembly was performed with MEGAHIT (v1.2.9) ^46^. Sequencing depth information for the assembled contigs was obtained using BWA (v0.7.17-r1198) ^47^, SAMtools (v1.18) ^48^, and the jgi_summarize_BAM_contig_depths script. Binning of the assembled contigs was carried out with MetaBAT2 (v2.15) ^49^, using the parameters “-m 2000 –s 200000 –-seed 2024.” The completeness and contamination of bins in the high-quality set were assessed with CheckM2 (v1.0.1) ^50^, retaining bins with ≥50% completeness and ≤10% contamination. Finally, metagenome-assembled genomes (MAGs) were dereplicated at 99% average nucleotide identity (ANI) using dRep (v3.4.3) ^51^ with the parameters “-pa 0.9 –sa 0.99.”

### Gene prediction and functional annotation

Taxonomic annotation of the MAGs was performed using GTDB-Tk (v2.3.2) ^52^. Open reading frames (ORFs) were predicted using Prodigal (v2.6.3) ^53^. Antibiotic resistance genes (ARGs) were identified by aligning protein sequences to the Comprehensive Antibiotic Resistance Database (CARD) ^54^ using DIAMOND (v2.1.8.162) ^55^, with sequence identity and query coverage both set to exceed 80%, and an e-value threshold of 1e-5. ARGs conferring resistance to at least two drug classes were classified as multi-drug resistant, while those resistant through at least two mechanisms were categorized as multi-type mechanism genes. Virulence factor genes (VFGs) were identified by aligning sequences to the Virulence Factor Database (VFDB) using DIAMOND (v2.1.8.162) ^55^, with sequence identity and query coverage set to exceed 80%. Mobile genetic elements (MGEs) were detected by aligning gene sequences to the Mobile Genetic Element Database using BLASTN (v2.13.0) with the parameters “-evalue 1e-5 –perc_identity 80 –qcov_hsp_perc 80.” To assess the abundance of ARGs, MGEs, and VFGs, 20 million clean reads per sample were mapped to reference genes using Bowtie2 (v2.5.0) ^45^ with default parameters. The read counts were normalized to transcripts per kilobase million (TPM).

### Correlation analysis

Correlation analysis was conducted using the R platform (v4.4.1). Procrustes analysis was performed with the procrustes function from the *vegan* package, and Mantel tests were conducted using the mantel_test function from the ‘LinkET’ package (v0.7.4). Spearman’s rank correlation coefficient was applied to separately assess diversity-based and abundance-based correlations between ARGs and virulence factor genes (VFGs), as well as between ARGs and mobile genetic elements (MGEs).

### Comparison of ARGs in gut microbiota of wild birds, chickens, and humans

We collected a total of 60,664 metagenome-assembled genomes (MAGs) from human gut microbiota (available at https://github.com/snayfach/IGGdb) and 26,053 MAGs from chicken gut microbiota. Specifically, the chicken-derived MAGs included 12,339 MAGs from the National Microbiology Data Center (NMDC, https://nmdc.cn/icrggc), 979 MAGs from the ENA database (project IDs: PRJEB55375 and PRJEB55374), and 6,786 MAGs from the Figshare database (https://dx.doi.org/10.6084/m9.figshare.24901884, https://dx.doi.org/10.6084/m9.figshare.24901878, https://doi.org/10.6084/m9.figshare.24681096.v1).

Additionally, 5,949 MAGs were retrieved from the NCBI database (project ID: PRJNA1099794). All MAGs were filtered using the same criteria as applied to wild bird samples: completeness ≥50% and contamination ≤10%. The filtered genomes were clustered at 99% average nucleotide identity (ANI), resulting in 36,467 human-derived and 17,013 chicken-derived gut microbiota MAGs. Subsequently, antibiotic resistance genes (ARGs) were annotated using the Comprehensive Antibiotic Resistance Database (CARD), and shared or host-specific ARG profiles were compared among the three host groups.

## Statistical analysis and visualization

The phylogenetic tree was visualized using the iTOL platform. Shannon and Simpson indices were calculated based on the relative abundance of functional genes. Principal Coordinate Analysis (PCoA) was applied to assess species diversity using Bray-Curtis distance, while Permutational Multivariate Analysis of Variance (PERMANOVA) was conducted with the ‘vegan’ (v2.6.10) and ‘dplyr’ (v1.1.4) packages to evaluate group differences and assess the independent and interactive effects of year, location, species, and residency status on ARG community structure. A total of 999 permutations were used to test statistical significance and quantify the proportion of explained variance. Gene arrow maps were generated using the ‘gggenes’ package (v0.5.1), and network graphs were visualized with the ‘ggraph’ package (v2.1.0). Genome structure and target gene annotation were performed using CGView services (https://proksee.ca/). Venn diagrams were created using the ‘ggvenn’ package (v0.1.10), while all other visualizations were generated with the ‘ggplot2’ package (v3.3.6). Statistical analyses were performed in R (v4.4.1).

## Ethical approval

This study was approved by the Institutional Animal Care and Use Committee of Qingdao Agricultural University (Approval no. QAU-AEW-20230107001).

## Declaration of competing interest

The authors declare that they have no known competing financial interests or personal relationships that could have influenced the work reported in this paper.

## Acknowledgements

The study was supported by the distinguished Scholar Research Fund of Qingdao Agricultural University (665-1120046).

## Data availability

The metagenomic sequencing data for the 50 wild bird gut microbiomes used in this study are available in the National Center for Biotechnology Information (NCBI) under accession number PRJNA1229141. The 10,455 gut microbial MAGs from wild birds generated in this study are available from the Figshare repository (https://doi.org/10.6084/m9.figshare.28504238). All other data supporting the findings of this study are provided in the paper and supplementary materials or can be obtained from the corresponding author(s) upon request.

## Author Contributions

**Jin-Wen Su:** Writing –original draft, Formal analysis, Software, Visualization. **Ying-Qian Gao:** Formal analysis, Software, Writing-review and editing. **Hany M. Elsheikha:** Conceptualization, Validation, Writing-review and editing. **Cong-Cong Lei:** Resources, Supervision, Visualization, Writing-review and editing. **Hong-Lin Xie:** Resources, Writing-review and editing. **Fu-Long Nan:** Validation, Supervision, Software, Writing-review and editing. **He Ma:** Supervision, Software, Writing-review and editing. **Meng-Ting Yang:** Validation, Writing-review and editing. **Hai-Tao Wang:** Validation, Writing-review and editing. **Hong-Bo Ni:** Conceptualization, Writing-review and editing. **He-Ting Sun:** Resources, Writing-review and editing. **Hong-Chao Sun:** Conceptualization, Formal analysis, Writing-review and editing. **Xiao-Xuan Zhang:** Conceptualization, Funding acquisition, Validation, Supervision, Software, Writing-review and editing.

## Supplementary Materials

### Additional file 1

**Supplementary Fig. 1:**
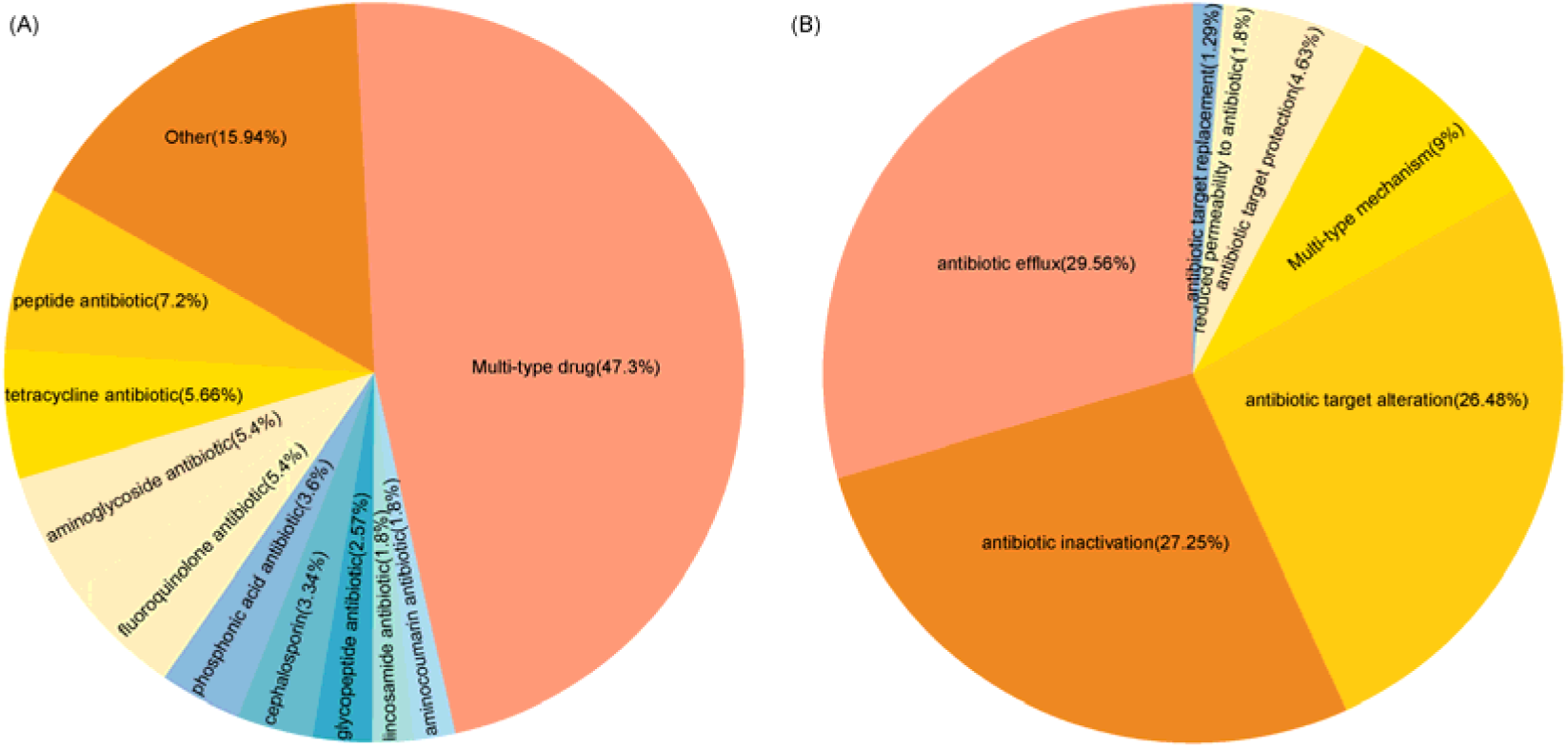
Distribution of resistance phenotypes and mechanisms. (A) Distribution of the ten most predominant resistance phenotypes. (B) Distribution of the ten most prevalent resistance mechanisms.

**Supplementary Fig. 2:**
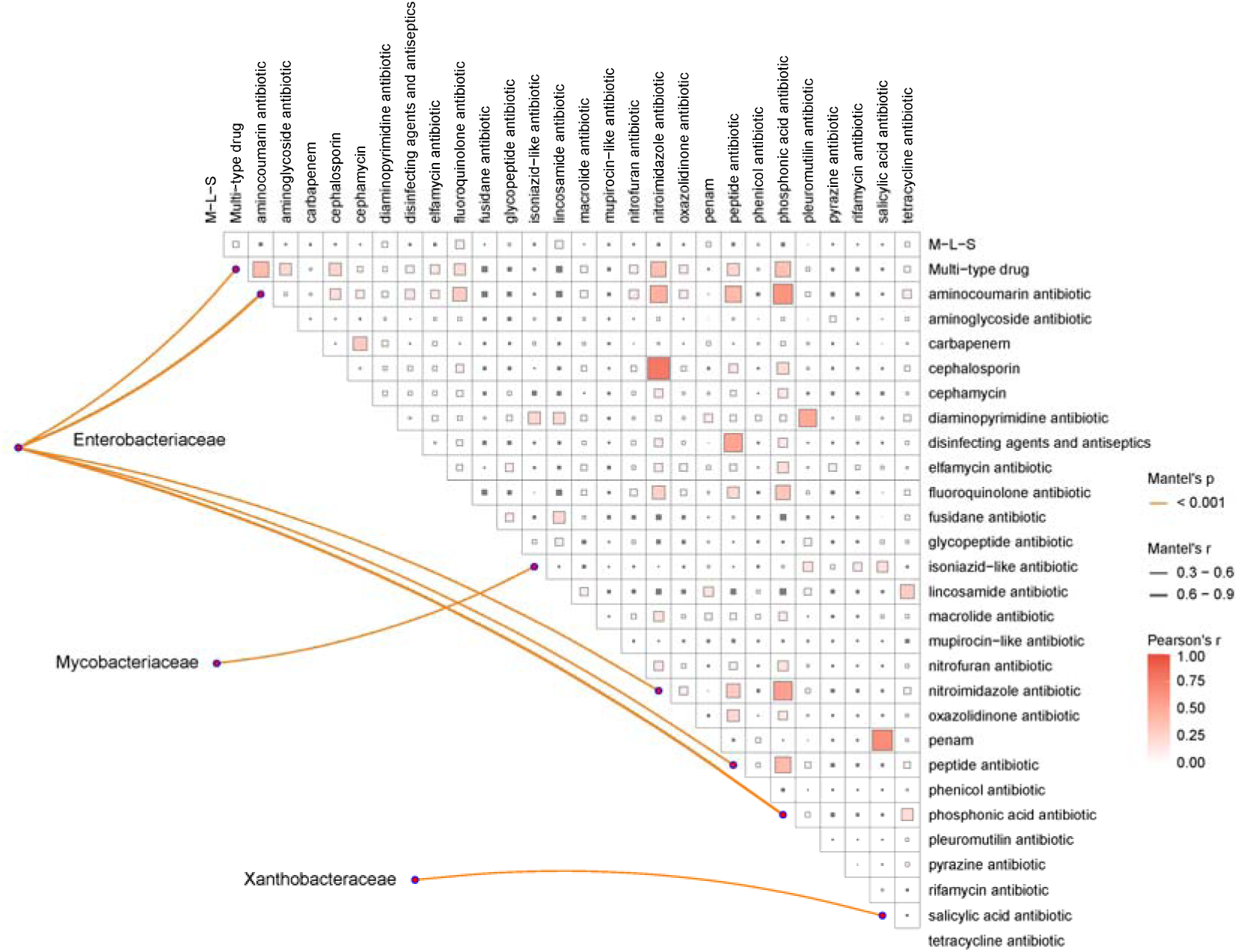
Mantel test analysis of microbial families and resistance phenotypes. Mantel test analysis of the correlation between the top 20 abundant microbial families and resistance phenotypes.

**Supplementary Fig. 3:**
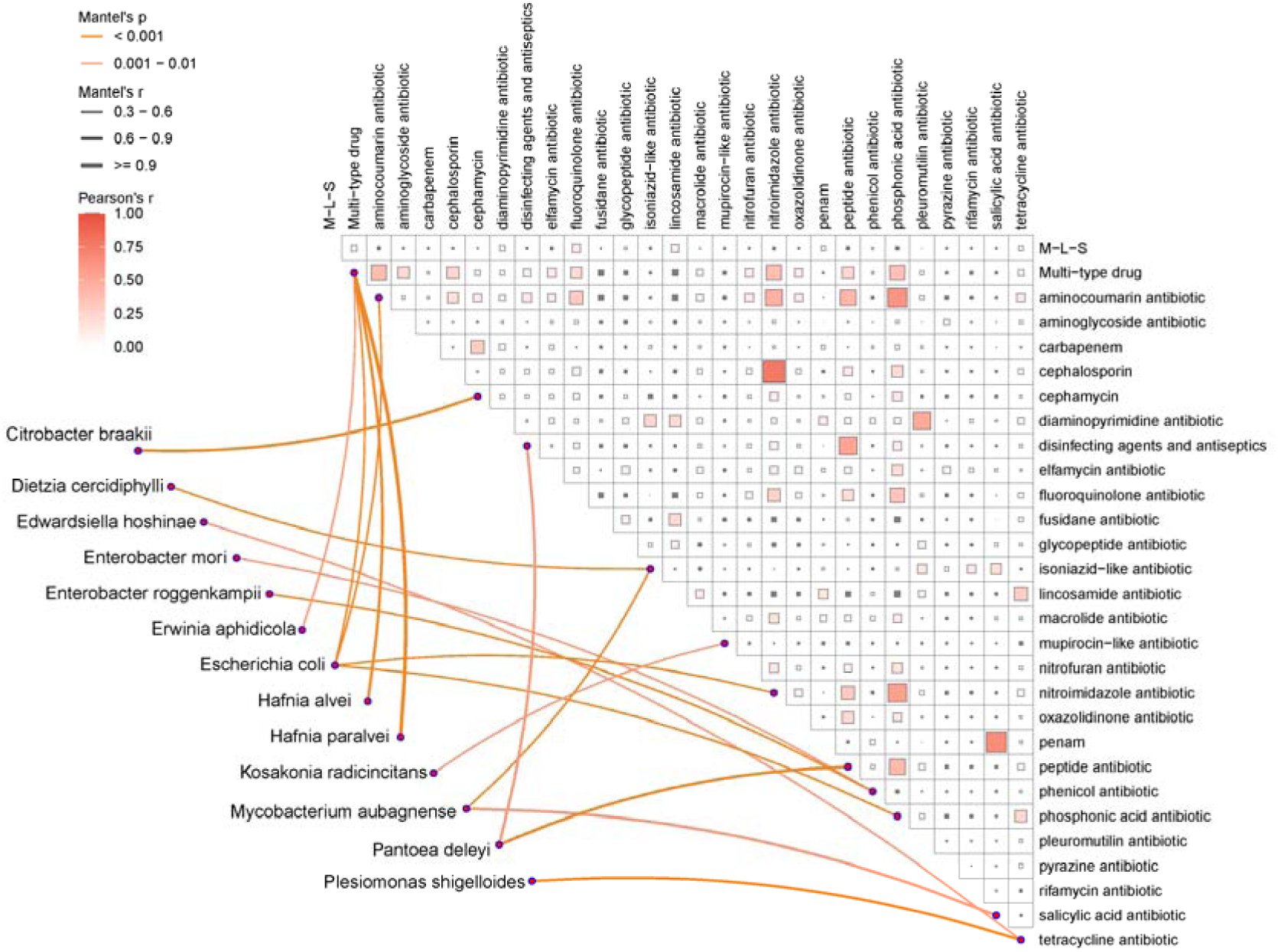
Mantel test analysis of microbial species and resistance phenotypes. Mantel test analysis of the correlation between species from three microbial families significantly associated with resistance phenotypes and the resistance phenotypes.

**Supplementary Fig. 4:**
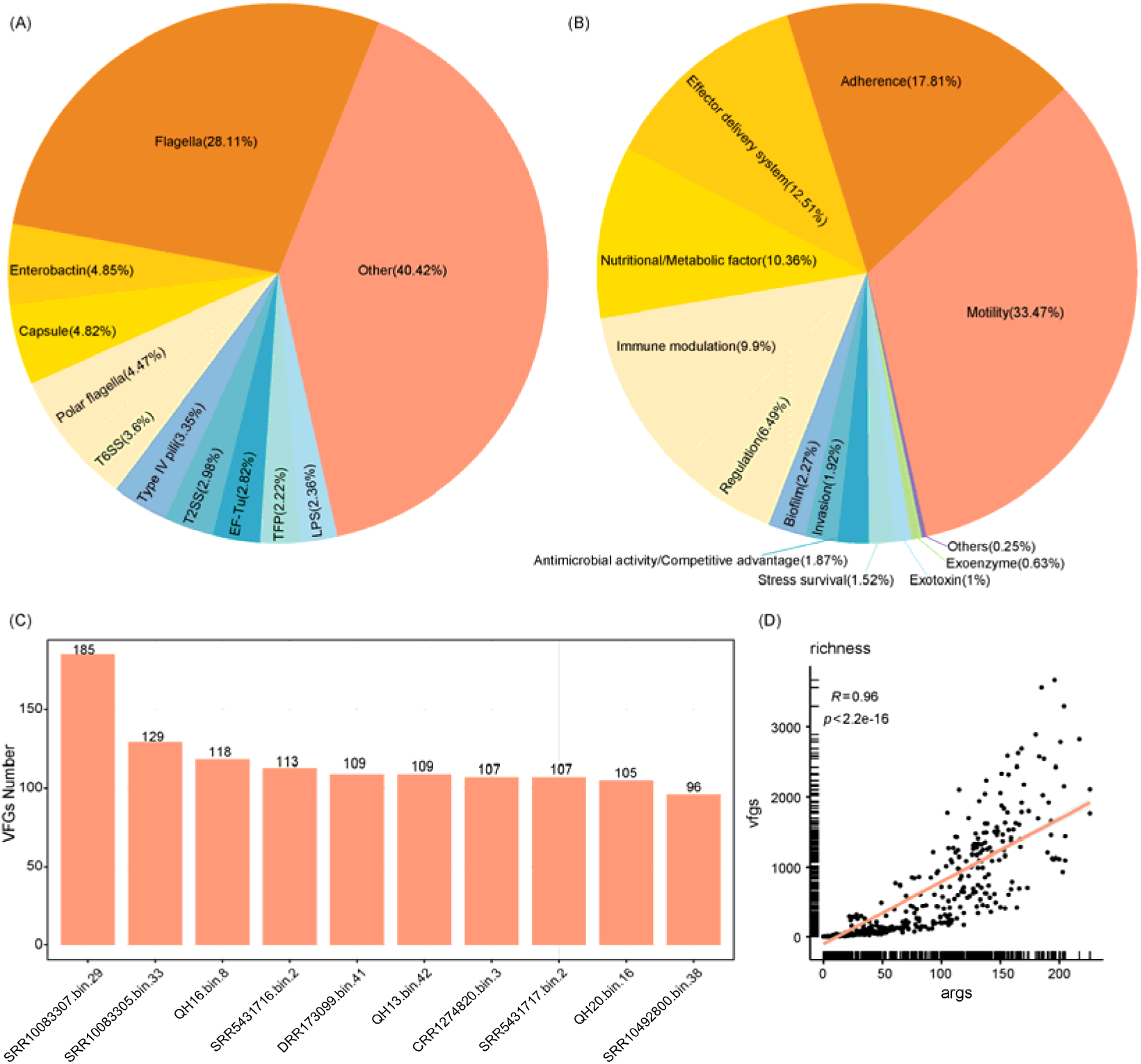
Distribution and correlation of VFs and VFGs. (A) Distribution of the ten most abundant VFs. (B) Distribution of the ten most abundant VFCs. (C) Top ten MAGs with the highest number of VFGs. (D) Correlation of the Richness index between ARGs and VFGs, calculated using Spearman’s rank correlation.

**Supplementary Fig. 5:**
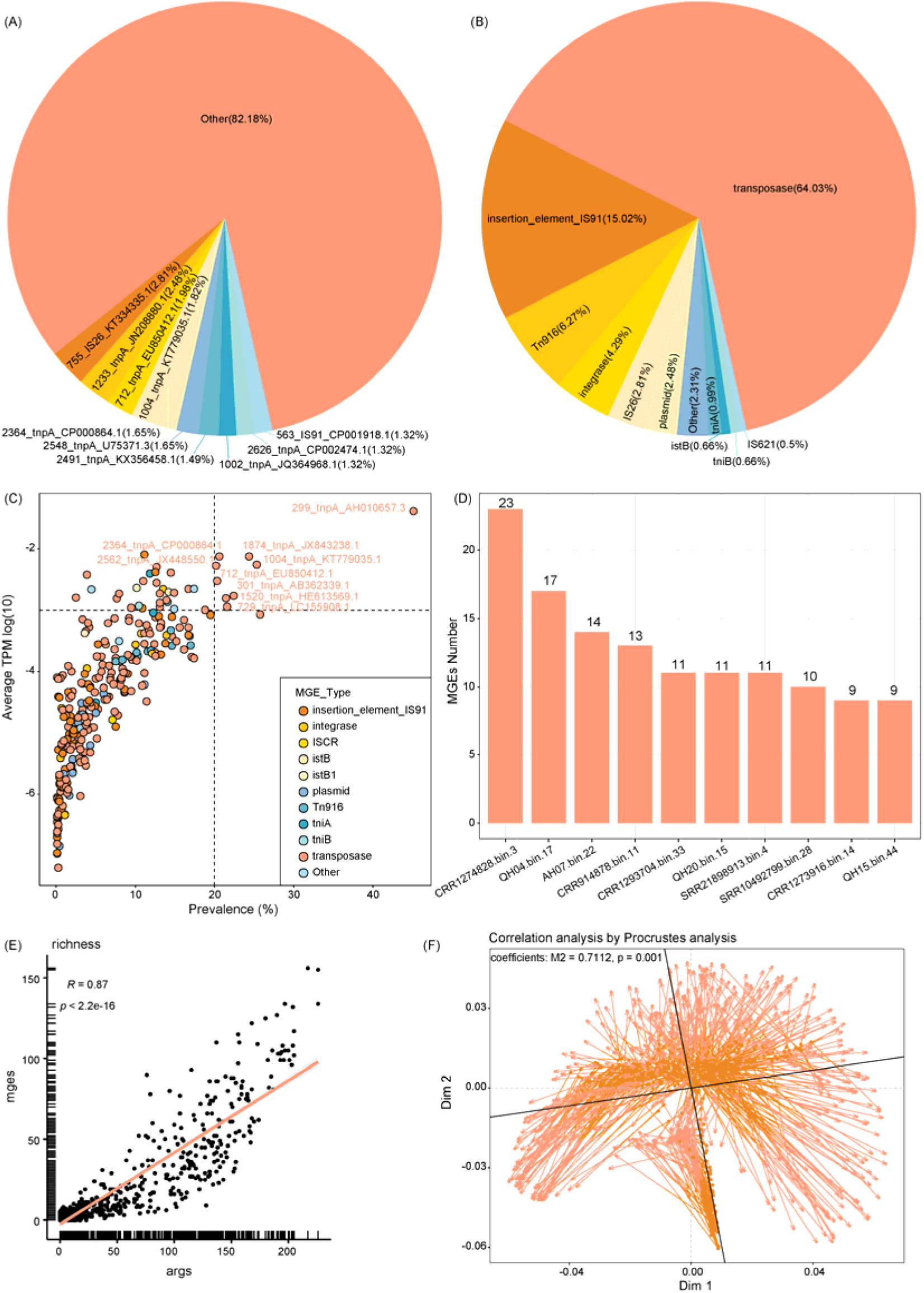
Distribution and correlation of MGEs and ARGs. (A) Distribution of the ten most abundant MGEs. (B) Distribution of the ten most abundant MGE types. (C) Prevalence of MGEs, with each dot representing an MGE and color indicating the corresponding MGE type. (D) Top ten MAGs with the highest number of MGEs. (E) Correlation of the Richness index between ARGs and MGEs, calculated using Spearman’s rank correlation. (F) Procrustes analysis showing a correlation between the abundance profiles of ARGs and MGEs.

**Supplementary Fig. 6:**
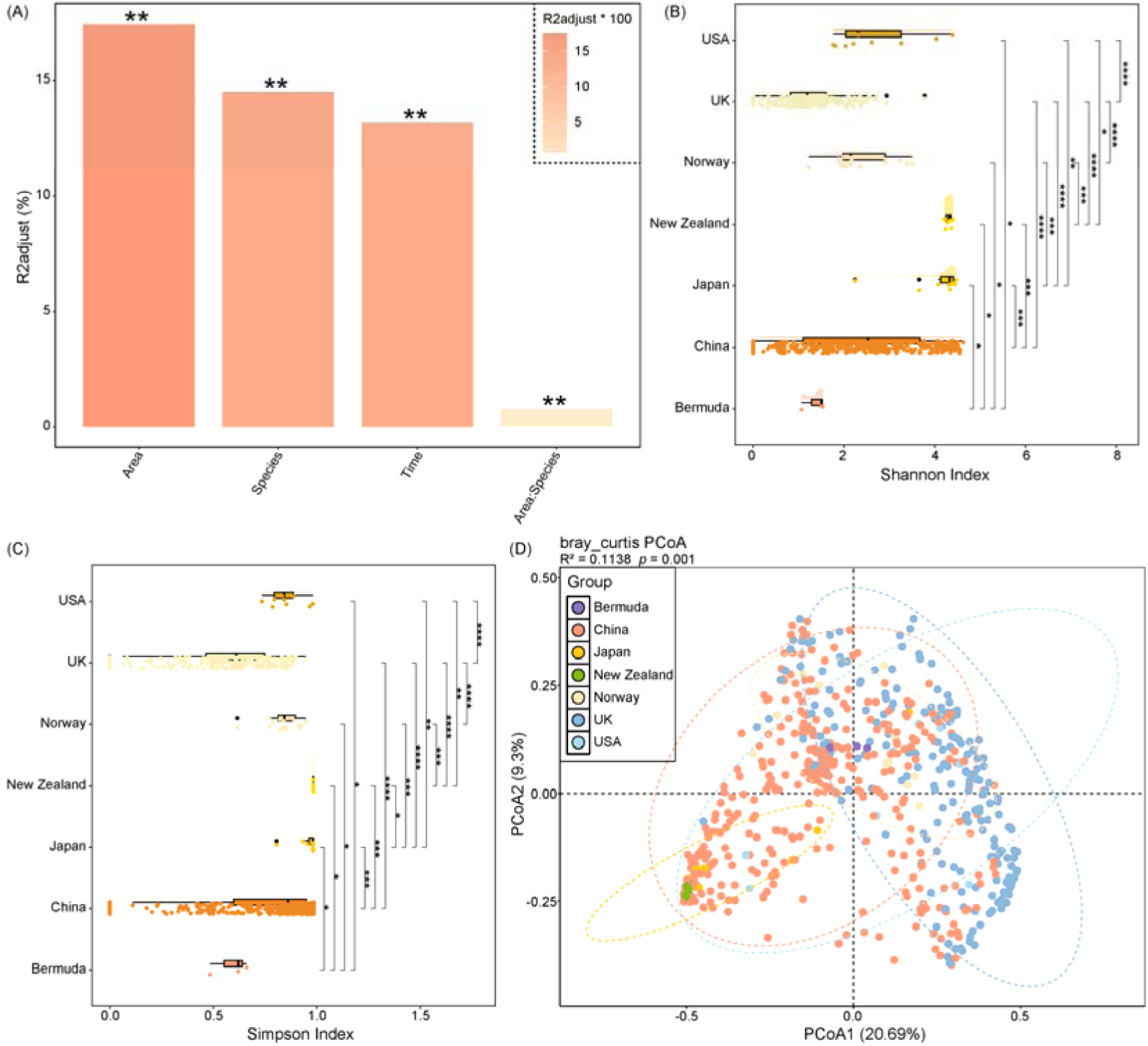
Analysis of factors influencing ARG composition. (A) PERMANOVA analysis based on Bray-Curtis distance matrix showing the effects of individual and interactive factors on the composition of ARGs in wild bird gut microbiota. The y-axis indicates adjusted R² values; ** denotes *p* < 0.01. (B-C) Boxplots showing Shannon and Simpson diversity indices of gut microbial ARGs across different sampling areas. Statistical significance was assessed using the Wilcoxon rank-sum test: * *p* < 0.05; ** *p* < 0.01; *** *p* < 0.001; **** *p* < 0.0001. (D) PCoA plot based on Bray-Curtis distances illustrating beta diversity differences in ARG composition among wild birds from different sampling areas. Points represent samples plotted along PCoA1 and PCoA2, with ellipses representing 95% confidence intervals.

**Supplementary Fig. 7:**
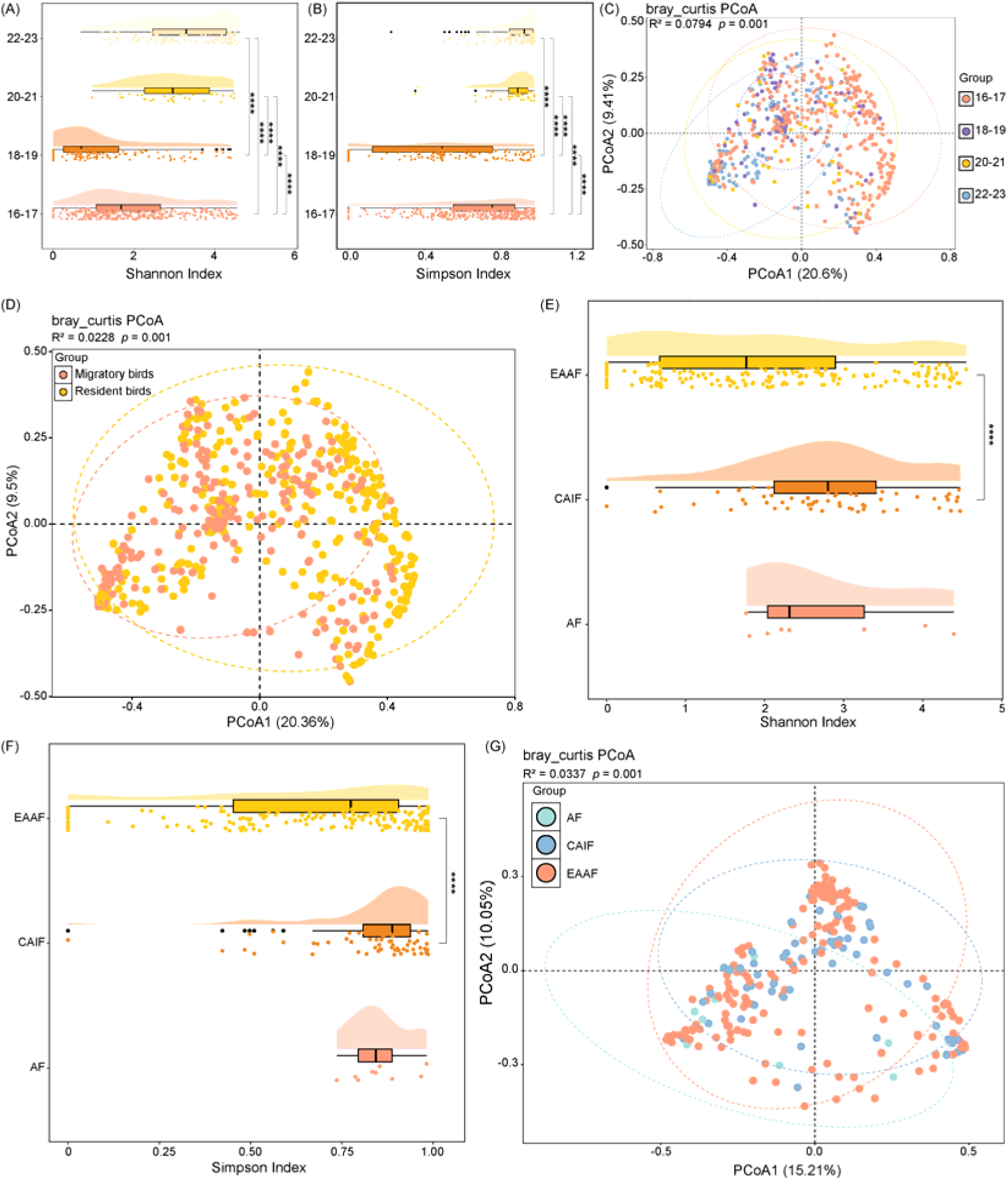
Analysis of ARG diversity across sampling times and bird migration. (A-B) Boxplots showing Shannon and Simpson diversity indices of gut microbial ARGs in wild birds across different sampling times. Statistical significance was assessed using the Wilcoxon rank-sum test: * *p* < 0.05; ** *p* < 0.01; *** *p* < 0.001; **** *p* < 0.0001. (C) PCoA plot based on Bray-Curtis distances illustrating beta diversity differences in gut microbial ARG composition of wild birds across sampling times. Points represent samples plotted along PCoA1 and PCoA2; ellipses indicate 95% confidence intervals for each group. (D) PCoA plot based on Bray-Curtis distances showing beta diversity differences in gut microbial ARG composition between migratory and resident birds. Points represent samples plotted along PCoA1 and PCoA2, with ellipses representing 95% confidence intervals. (E-F) Boxplots showing Shannon and Simpson diversity indices of gut microbial ARGs in wild birds across different migratory flyways. Statistical significance was assessed using the Wilcoxon rank-sum test: * *p* < 0.05; ** *p* < 0.01; *** *p* < 0.001; **** *p* < 0.0001. (G) PCoA plot based on Bray-Curtis distances showing beta diversity differences in gut microbial ARG composition among wild birds from different migratory flyways. Points represent samples plotted along PCoA1 and PCoA2, and ellipses represent 95% confidence intervals for each group.

**Supplementary Fig. 8:**
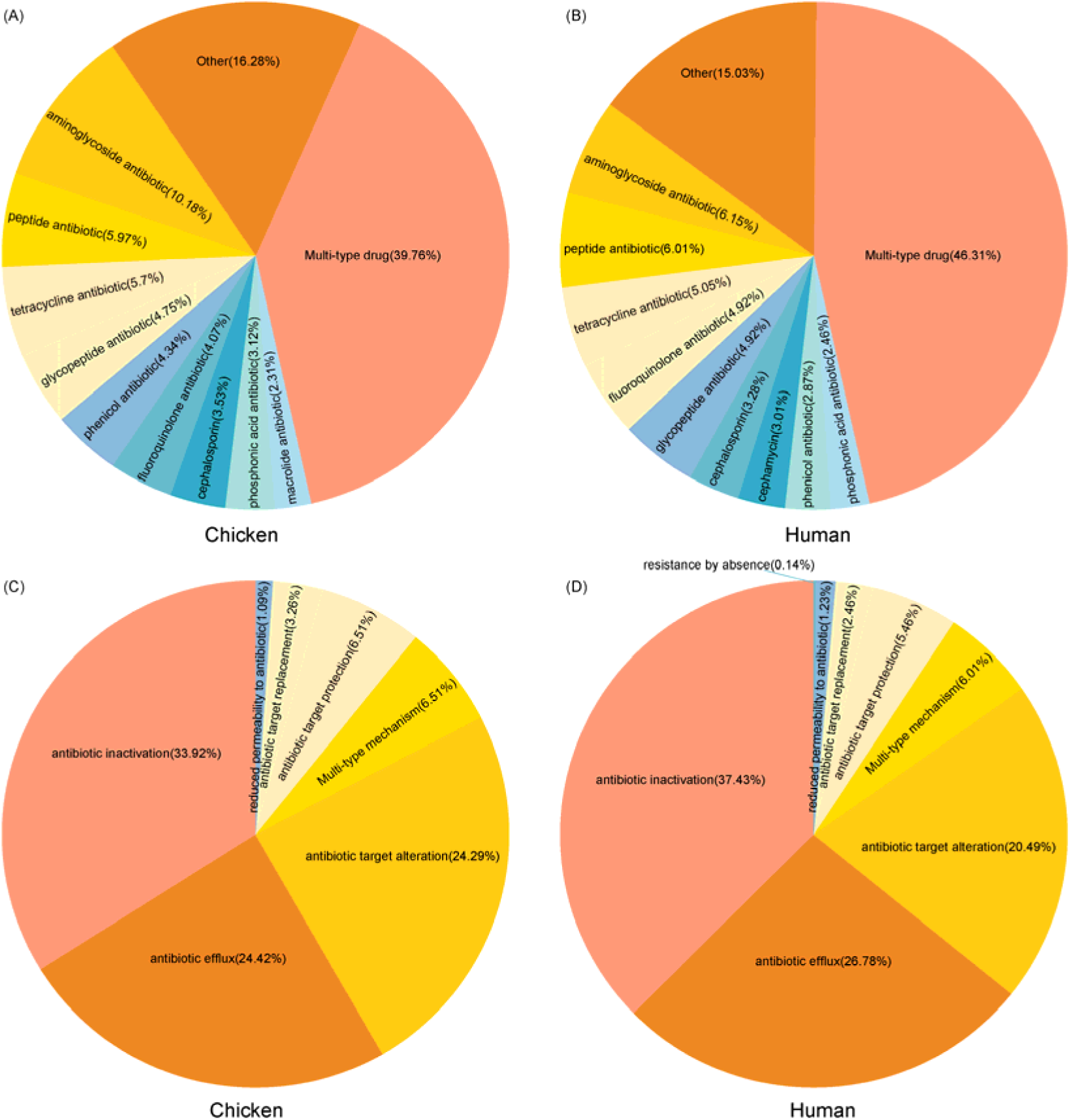
Comparison of ARG types and resistance mechanisms in chickens, humans, and wild birds. (A) Top ten ARG types in chicken gut microbiota. (B) Top ten ARG types in human gut microbiota. (C) Top ten resistance mechanisms of ARGs in chicken gut microbiota. (D) Top ten resistance mechanisms of ARGs in human gut microbiota.

### Additional file 2

**Supplementary Table 1:** Overview of the 718 analyzed samples.

**Supplementary Table 2:** Characteristics of 2,516 metagenome-assembled genomes

**Supplementary Table 3:** Annotation details of 5,596 antibiotic resistance genes.

**Supplementary Table 4:** Prevalence data of identified ARGs.

**Supplementary Table 5:** Annotation details of 8,090 virulence factor genes.

**Supplementary Table 6:** Prevalence data of identified VFGs.

**Supplementary Table 7:** Significant correlations between the top 20 most prevalent VFGs and ARGs.

**Supplementary Table 8:** Annotation details of 606 MGE genes.

**Supplementary Table 9:** Prevalence data of identified MGEs.

**Supplementary Table 10:** Significant correlations between the top 20 most prevalent MGEs and ARGs.

**Supplementary Table 11:** Shannon and Simpson diversity indices of ARGs across various species.

**Supplementary Table 12:** ARG prevalence across different migratory bird routes.

**Supplementary Table 13:** Shared ARGs between the gut microbiota of wild birds and humans.

**Supplementary Table 14:** Compilation of common ARGs conferring resistance to tigecycline, vancomycin, polymyxins, and β-lactams.

## Notes

### Competing Interest Statement

The authors have declared no competing interest.

