## Additional file 1 for "Integrated metagenome-resolved profiling of the resistome, virulome, and mobilome in the gut microbiota of wild birds"

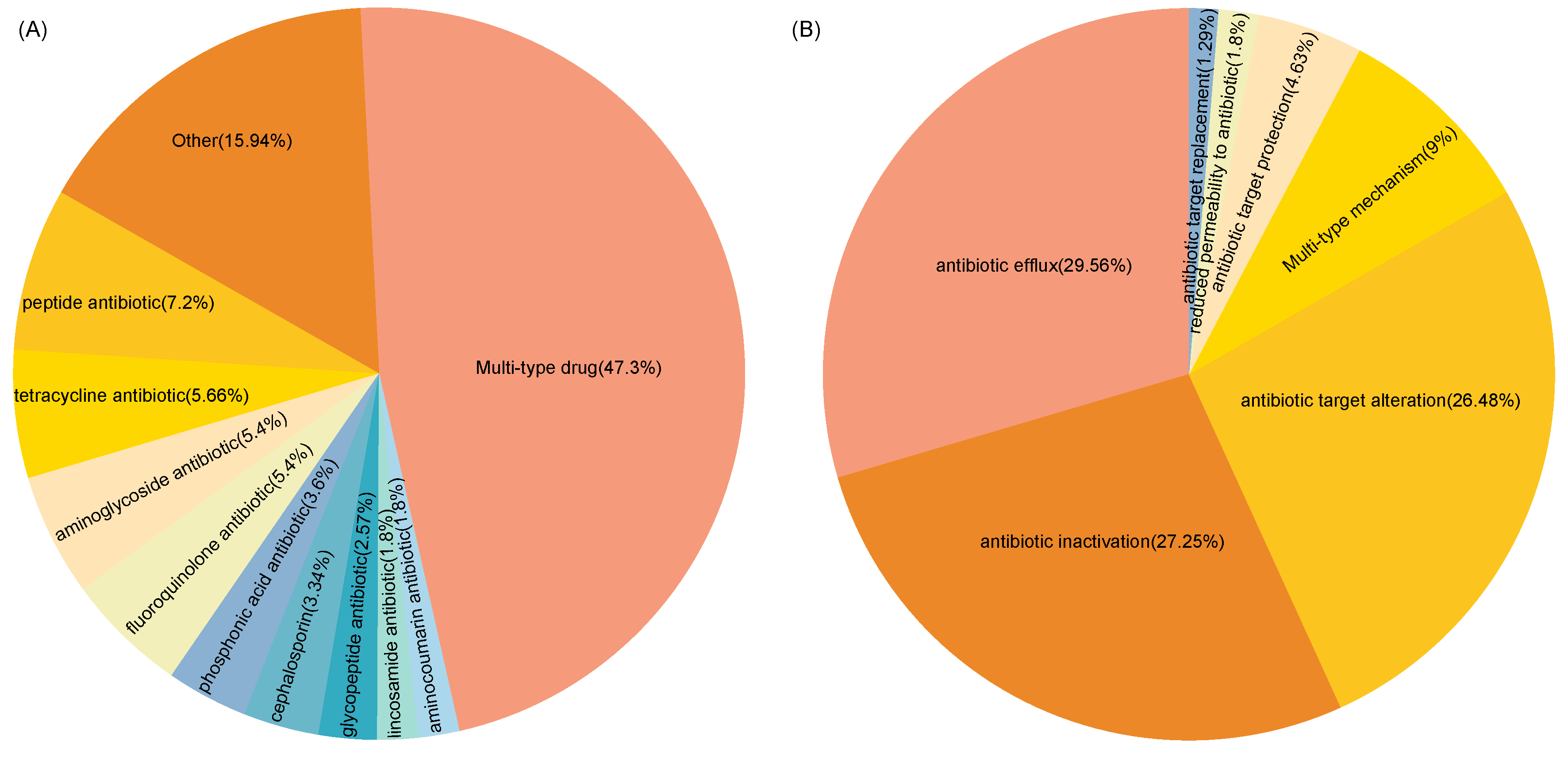

**Supplementary Fig. 1: Distribution of resistance phenotypes and mechanisms** (A) Distribution of the ten most predominant resistance phenotypes. (B) Distribution of the ten most prevalent resistance mechanisms.

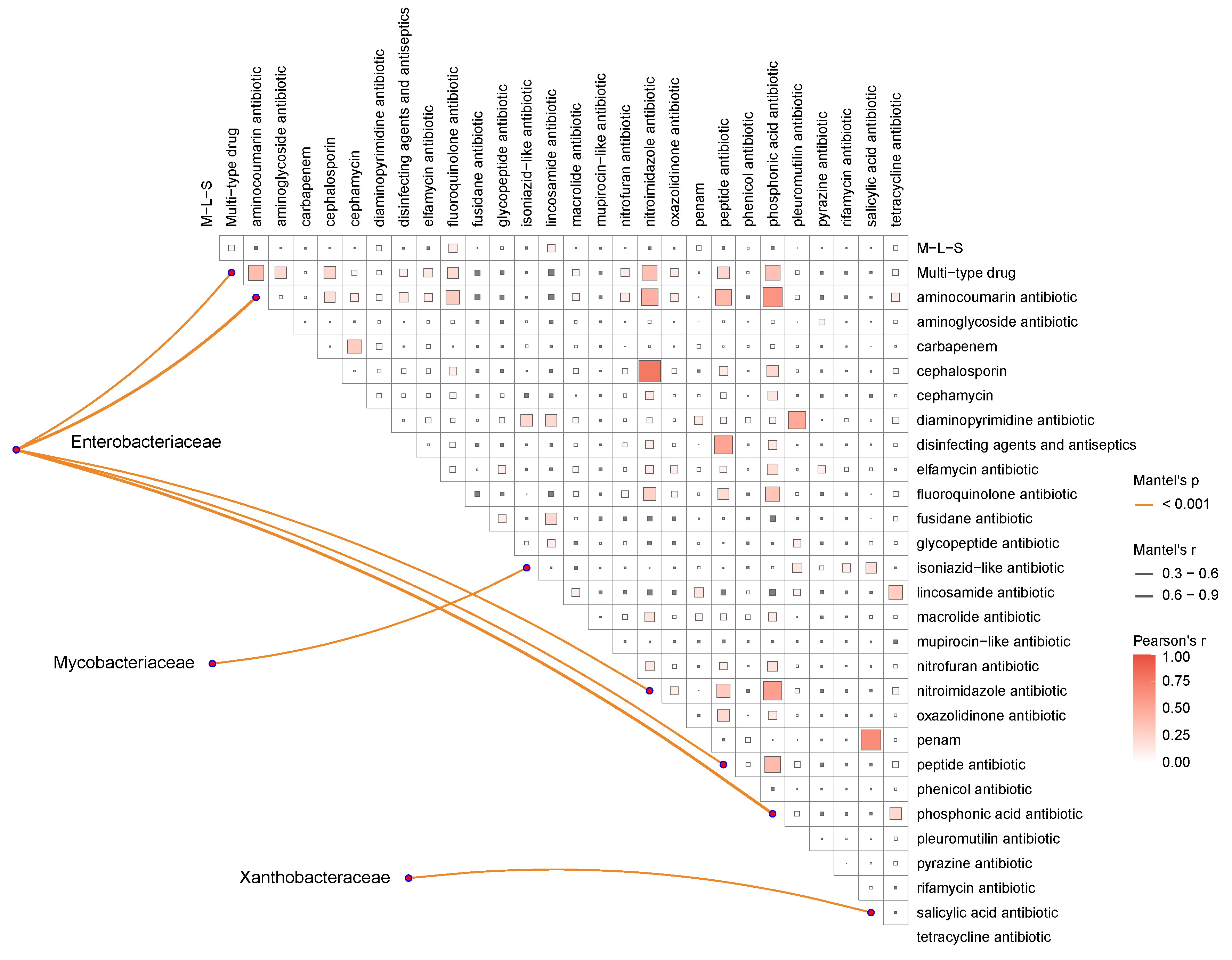

**Supplementary Fig. 2:** **Mantel test analysis of microbial families and resistance phenotypes.** Mantel test analysis of the correlation between the top 20 abundant microbial families and resistance phenotypes.

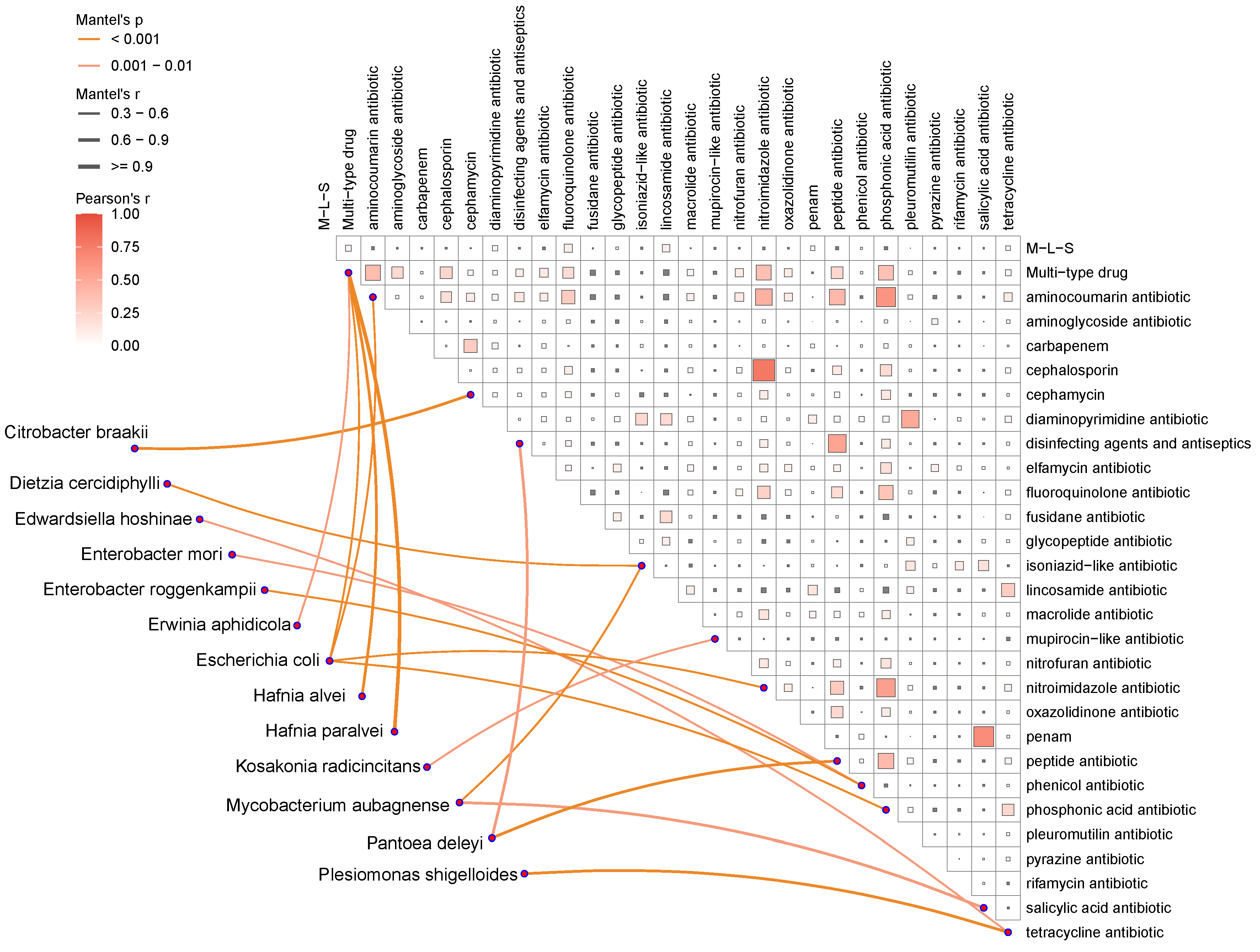

**Supplementary Fig. 3:** **Mantel test analysis of microbial species and resistance phenotypes.** Mantel test analysis of the correlation between species from three microbial families significantly associated with resistance phenotypes and the resistance phenotypes.

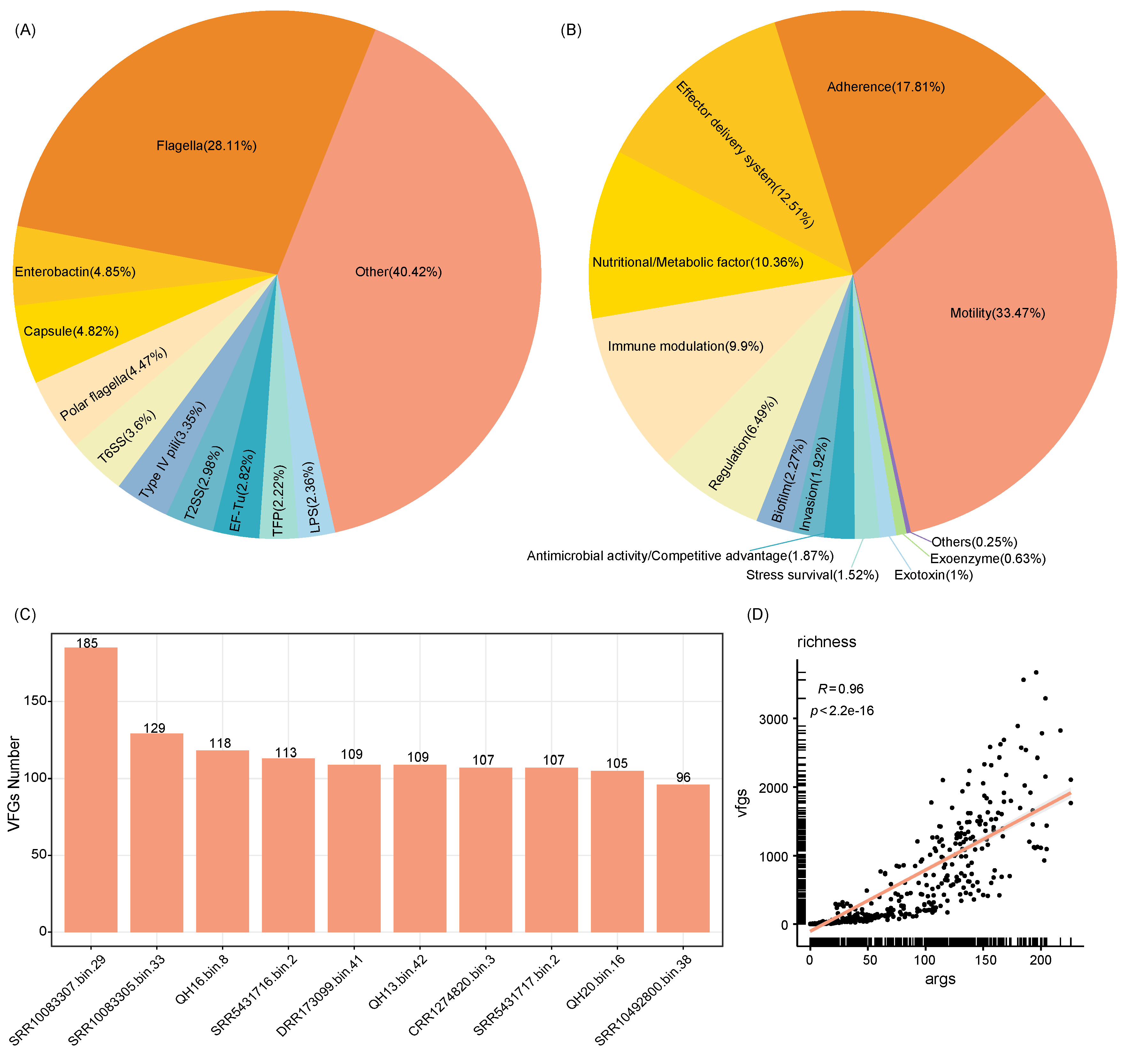

**Supplementary Fig. 4: Distribution and correlation of VFs and VFGs.** (A) Distribution of the ten most abundant VFs. (B) Distribution of the ten most abundant VFCs. (C) Top ten MAGs with the highest number of VFGs. (D) Correlation of the Richness index between ARGs and VFGs, calculated using Spearman’s rank correlation.

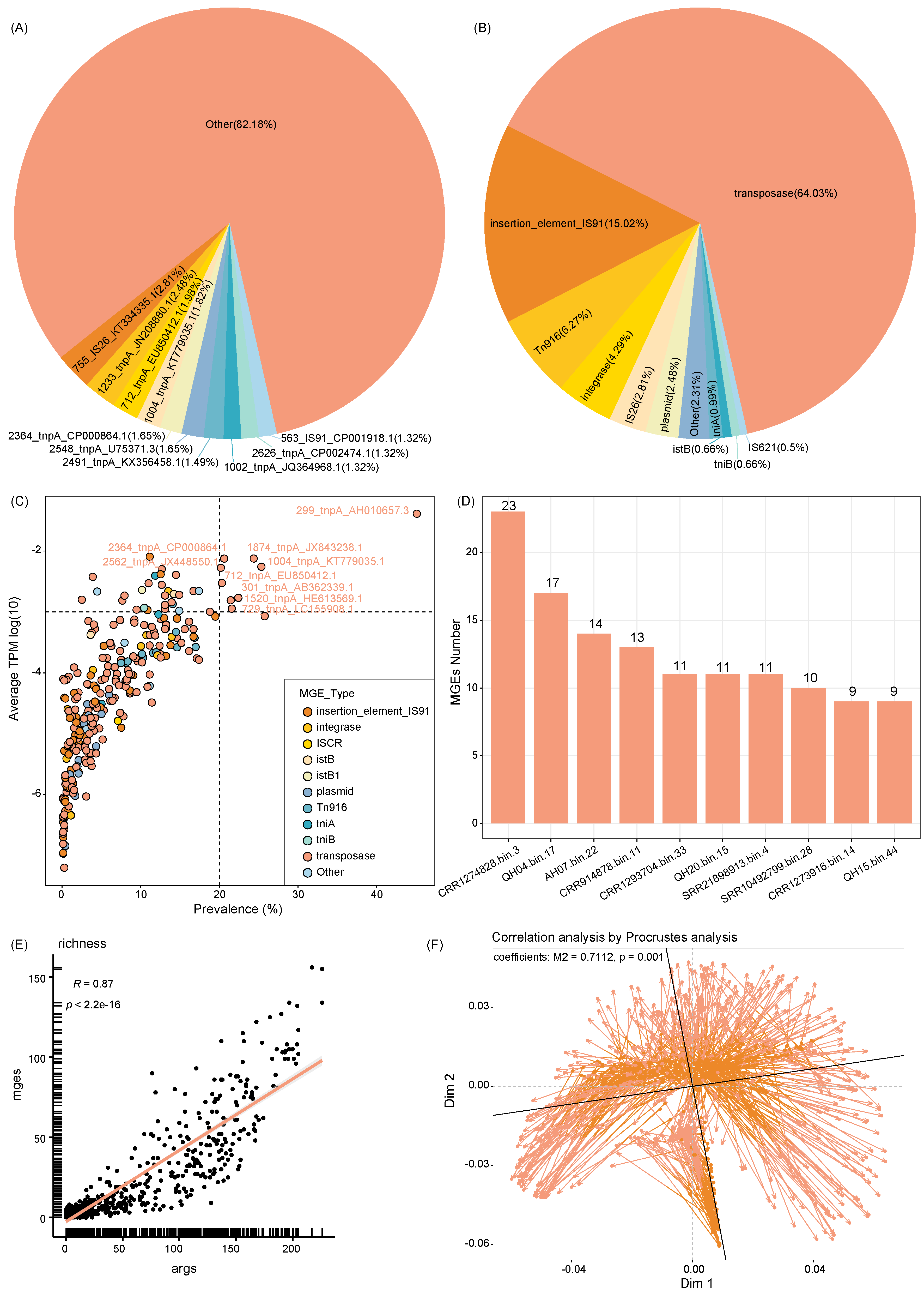

**Supplementary Fig. 5: Distribution and correlation of MGEs and ARGs.** (A) Distribution of the ten most abundant MGEs. (B) Distribution of the ten most abundant MGE types. (C) Prevalence of MGEs, with each dot representing an MGE and color indicating the corresponding MGE type. (D) Top ten MAGs with the highest number of MGEs. (E) Correlation of the Richness index between ARGs and MGEs, calculated using Spearman’s rank correlation. (F) Procrustes analysis showing a correlation between the abundance profiles of ARGs and MGEs.

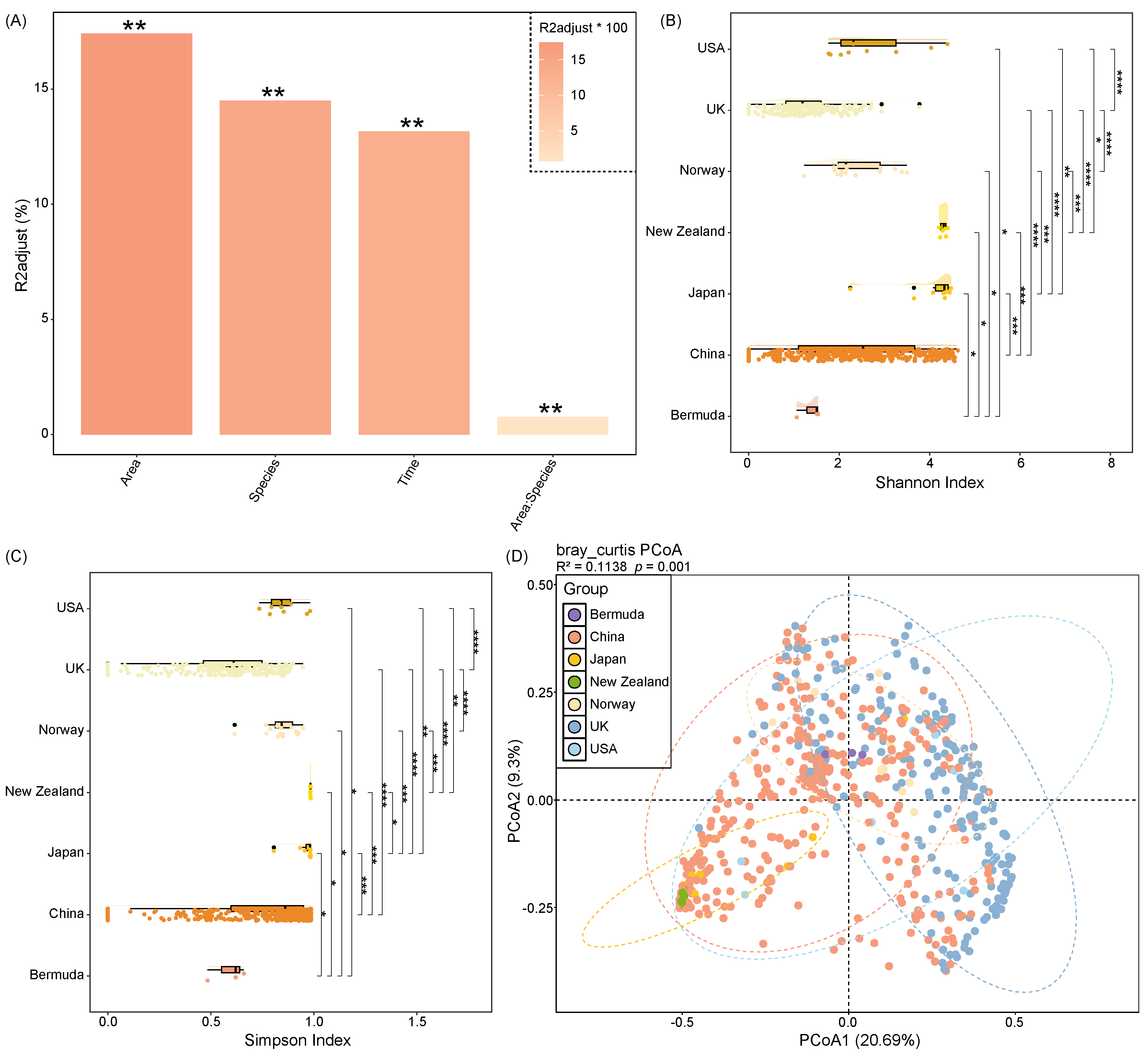

**Supplementary Fig. 6: Analysis of factors influencing ARG composition.** (A) PERMANOVA analysis based on Bray-Curtis distance matrix showing the effects of individual and interactive factors on the composition of ARGs in wild bird gut microbiota. The y-axis indicates adjusted R² values; ** denotes *p* < 0.01. (B-C) Boxplots showing Shannon and Simpson diversity indices of gut microbial ARGs across different sampling areas. Statistical significance was assessed using the Wilcoxon rank-sum test: * *p* < 0.05; ** *p* < 0.01; *** *p* < 0.001; **** *p* < 0.0001. (D) PCoA plot based on Bray-Curtis distances illustrating beta diversity differences in ARG composition among wild birds from different sampling areas. Points represent samples plotted along PCoA1 and PCoA2, with ellipses representing 95% confidence intervals.

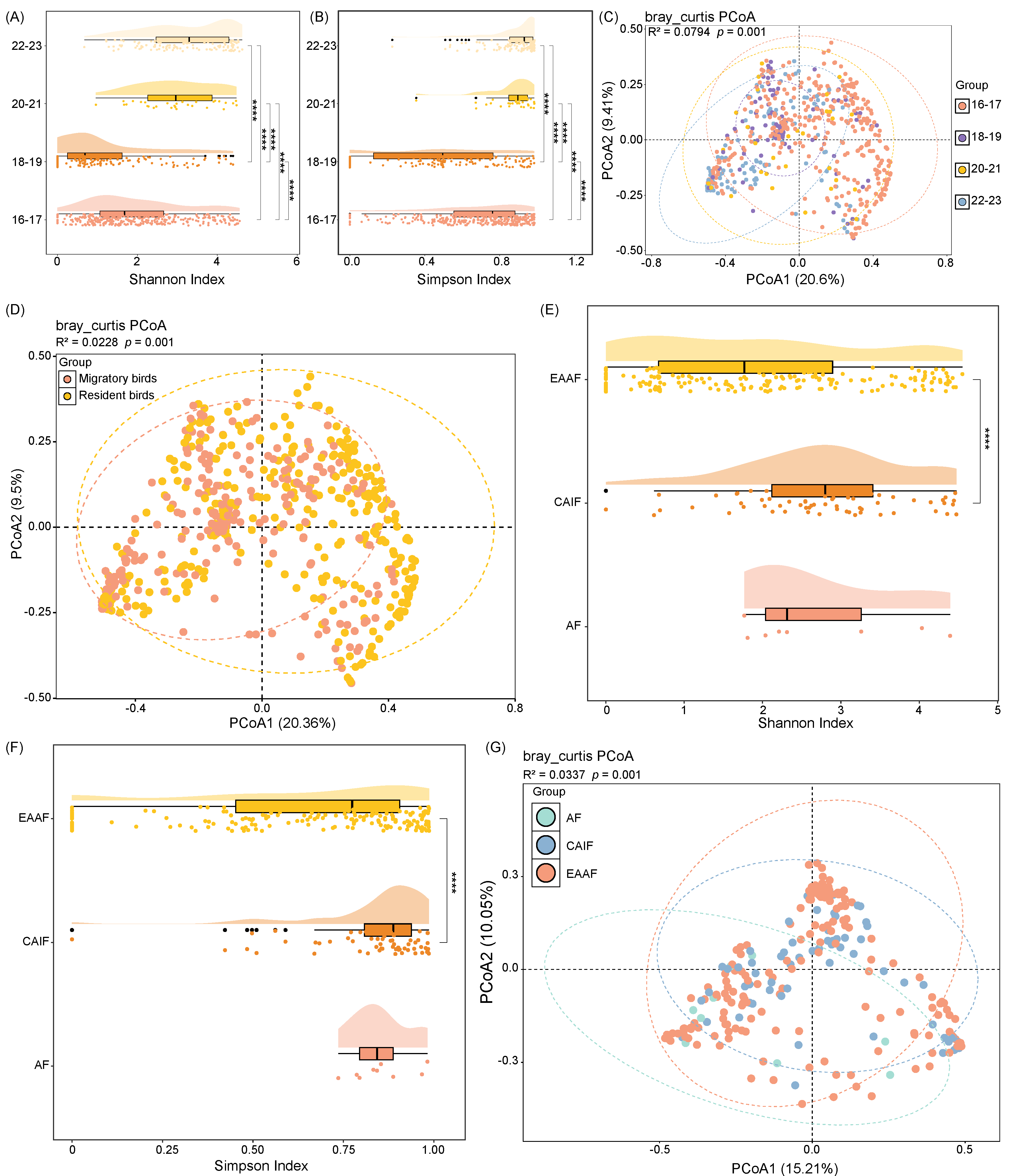

**Supplementary Fig. 7: Analysis of ARG diversity across sampling times and bird migration.** (A-B) Boxplots showing Shannon and Simpson diversity indices of gut microbial ARGs in wild birds across different sampling times. Statistical significance was assessed using the Wilcoxon rank-sum test: * *p* < 0.05; ** *p* < 0.01; *** *p* < 0.001; **** *p* < 0.0001. (C) PCoA plot based on Bray-Curtis distances illustrating beta diversity differences in gut microbial ARG composition of wild birds across sampling times. Points represent samples plotted along PCoA1 and PCoA2; ellipses indicate 95% confidence intervals for each group. (D) PCoA plot based on Bray-Curtis distances showing beta diversity differences in gut microbial ARG composition between migratory and resident birds. Points represent samples plotted along PCoA1 and PCoA2, with ellipses representing 95% confidence intervals. (E-F) Boxplots showing Shannon and Simpson diversity indices of gut microbial ARGs in wild birds across different migratory flyways. Statistical significance was assessed using the Wilcoxon rank-sum test: * *p* < 0.05; ** *p* < 0.01; *** *p* < 0.001; **** *p* < 0.0001. (G) PCoA plot based on Bray-Curtis distances showing beta diversity differences in gut microbial ARG composition among wild birds from different migratory flyways. Points represent samples plotted along PCoA1 and PCoA2, and ellipses represent 95% confidence intervals for each group.

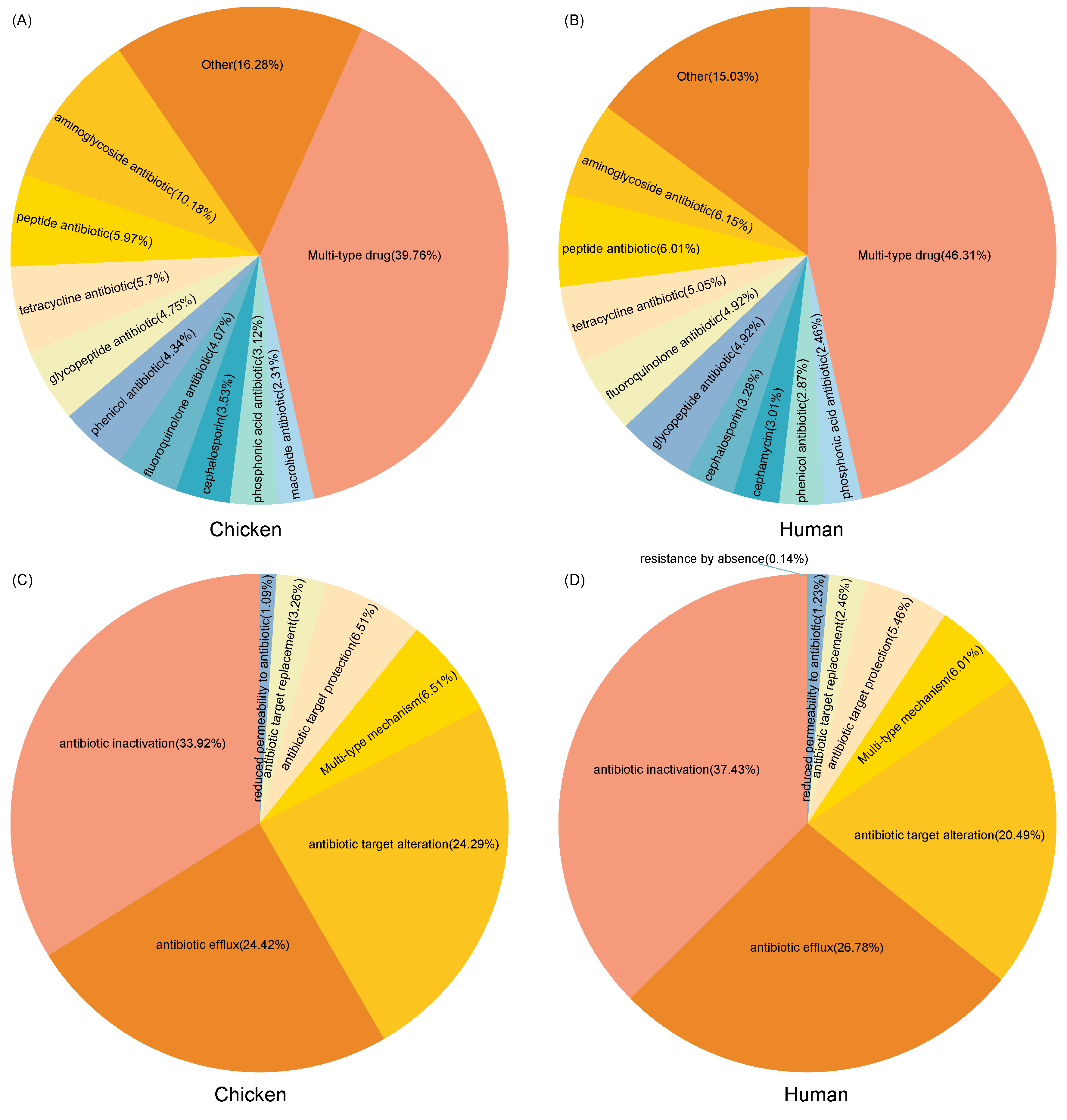

**Supplementary Fig. 8: Comparison of ARG types and resistance mechanisms in chickens, humans, and wild birds.** (A) Top ten ARG types in chicken gut microbiota. (B) Top ten ARG types in human gut microbiota. (C) Top ten resistance mechanisms of ARGs in chicken gut microbiota. (D) Top ten resistance mechanisms of ARGs in human gut microbiota.
